# Morphological Landscapes from High Content Imaging Identify Optimal Priming Strategies that Enhance MSC Immunosuppression

**DOI:** 10.1101/2021.02.23.432501

**Authors:** Seth H. Andrews, Matthew W. Klinker, Steven R. Bauer, Ross A. Marklein

## Abstract

Successful clinical translation of mesenchymal stromal cell (MSC) products has not been achieved in the United States and may be in large part due to MSC functional heterogeneity. Efforts have been made to identify ‘priming’ conditions that produce MSCs with consistent immunomodulatory function; however, challenges remain with predicting and understanding how priming impacts MSC behavior. The purpose of this study was to develop a high throughput, image-based approach to assess MSC morphology in response to combinatorial priming treatments and establish morphological profiling as an effective approach to screen the effect of manufacturing changes (i.e. priming) on MSC immunomodulation. We characterized the morphological response of multiple MSC lines/passages to an array of Interferon-gamma (IFN-γ) and Tumor Necrosis Factor alpha (TNF-α) priming conditions, as well as the effects of priming on MSC modulation of activated T cells and MSC secretome. Although considerable functional heterogeneity, in terms of T cell suppression, was observed between different MSC lines and at different passages, this heterogeneity was significantly reduced with combined IFN-γ/TNF-α priming. The magnitude of this change correlated strongly with multiple morphological features and was also reflected by MSC secretion of immunomodulatory factors e.g. PGE2, ICAM-1, and CXCL16. Overall, this study further demonstrates the ability of priming to enhance MSC function, as well as the ability of morphology to better understand MSC heterogeneity and predict changes in function due to manufacturing.

## 1. INTRODUCTION

Mesenchymal stromal cells (MSCs) are widely studied as potential treatments for immune-mediated diseases, such as osteoarthritis, diabetes, multiple sclerosis, and Alzheimer’s Disease^1–3^ via their modulation of immune cells such as T cells, B cells, macrophages, microglia, and dendritic cells.^1,4,5^ This immunomodulation has been shown to be mediated *in vitro* through the release of a multitude of immunomodulatory factors (termed ‘secretome’).^6^ MSCs can be used in an allogeneic, off-the-shelf manner, and have been studied in over 600 clinical trials with most studies reporting a good safety profile^7^; however, the manufacture of MSCs with consistent and predictable quality has proven difficult.^8^ This is due to rare employment of robust quantitative assays to assess their function, a lack of standardization in manufacturing processes, as well as MSC heterogeneity.^9,10^ MSC functional heterogeneity can be attributed to a number of factors, including donor/tissue sources, extended culture, and varying manufacturing processes.^10^

Although efforts have been made to develop improved assays for assessing and predicting MSC function^3,11,12^, the majority of studies still predominantly characterize MSCs using the ISCT criteria established in 2006.^13^ As the current prevailing hypothesis of MSC mechanism of action has centered on their immunomodulatory capacity, new assays have been proposed that are more functionally relevant in the context of immune diseases as compared to, for example, trilineage differentiation potential. For example, a number of immunomodulatory factors (such as IDO and PD-L1)^14,15^ have been demonstrated to be predictive of MSC suppression of activated T cells, which is one of the most common *in vitro* assays for MSC immunomodulatory activity. However, these assays are typically performed at a population level and cannot determine differences in MSC heterogeneity that may be present at the single cell level. Furthermore, single metrics may not fully capture the complex, multifactorial mechanisms of MSC immunomodulation in terms of their secretome and the diversity of target immune cells, and therefore multiple assays may be required to sufficiently capture MSC immunomodulatory capacity.^11,15^ Finally, the mechanisms of action of MSCs are still being explored, adding further challenge to the development of characterization assays.^8^

MSC heterogeneity also exists in terms of cell morphology, and MSC morphology has been shown to be predictive of several therapeutically relevant functions (osteogenesis^16^, chondrogenesis^17^, and immunosuppression^18,19^). As a characterization assay for MSCs, morphological profiling offers several advantages. First, it can be performed in a rapid, low-cost, and high-throughput manner^20,21^. Additionally, morphology is assessed at single-cell resolution, which is essential in recognizing and addressing MSC heterogeneity. Finally, cell morphology can represent a summation of complex signaling pathways,^22–24^ possibly serving as more effective critical quality attributes (CQAs i.e. predictors of quality) than expression of a single protein or gene.

No standardized approaches exist for manufacturing MSCs, therefore differences in manufacturing conditions (e.g. culture medium, vessels, cryopreservation) can significantly contribute to functional heterogeneity.^25^ An increasingly used strategy to mitigate heterogeneity is to prime (i.e. precondition, pretreat) MSCs in the presence of different microenvironmental signals to improve their immunomodulatory capacity.^26^ Examples of these priming signals include inflammatory cytokines such as IFN-γ and TNF-α, hypoxia, and certain biomaterials and 3D cultures.^27^ The effects of priming MSCs with inflammatory cytokines has been extensively studied, and can be assessed based on expression of factors such as IDO.^28,29^ However, to date studies examining IFN-γ and TNF-α priming have often been done in a low throughput, binary manner^26,30,31^, limiting our understanding of the interplay of different cytokines and dosing on MSC behavior.

The objective of this study was to develop and apply a high throughput morphological screening approach to identify optimal priming conditions that enhance MSC immunomodulatory function *in vitro*. We accomplished this by comprehensively profiling MSC morphological responses to a combinatorial array of cytokine priming conditions (IFN-γ/TNF-α) using high content imaging and automated image analysis of single cell morphological features (akin to image-based drug screens). IFN-γ/TNF-α priming ‘hits’ from morphological profiling were then assayed for their T cell suppressive function to identify priming conditions with enhanced immunosuppression as compared to unprimed MSCs. Finally, we examined MSC secretion of proteins relevant to immune activity and correlated those with T cell suppression. We identified multiple MSC morphological features that predicted the effects of IFN-γ and TNF-α priming in terms of enhanced T cell modulation. Additionally, these changes in morphology and function due to priming were reflected by MSC secretion of cytokines and chemokines associated with T cell activation. This further establishes morphology as a predictor of MSC function, as well as presents a generalizable strategy for screening manufacturing conditions to improve the function of promising cell therapies.

## 2. METHODS

### 2.1 MSC Manufacturing

Human bone marrow-derived MSCs were obtained from 4 different donors purchased from Lonza (Walkersville, MD, USA), AllCells (Emeryville, CA, USA), and RoosterBio (Frederick, MD, USA) (see **Table S1** for donor information). MSC culture conditions for the Lonza and AllCells (8F3560, 110877, PCBM1662) were chosen based on well-established protocols.^32^ Briefly, MSCs were continuously expanded in complete MSC growth medium (GM) containing 10% FBS (JMBiosciences), 1% L-glutamine (Invitrogen), and 1% penicillin/streptomycin in alpha-MEM (both Invitrogen) at a seeding density of 60 cells/cm^2^ in T-175 flasks for a total of 7 passages with MSCs cryopreserved at passages 3, 5, and 7. Passage 3 (P3) and passage 7 (P7) MSCs from donors 8F3560, 110877, and PCBM1662 were used in this study. The RoosterBio cell-line RB9 was expanded using RoosterBio’s recommended protocol, which consisted of seeding 10×10^6^ MSCs in 12 T-225 flasks (3,704 cells/cm^2^) in RoosterBio growth medium and culturing until 80-90% confluency. RB9 MSCs were continuously expanded with a portion of the harvested cells at each passage cryopreserved to create a cell bank with passage 2 and passage 5 RB9 MSCs used in this study. All MSC lines used in this study have been extensively characterized for their surface marker expression, genomic, epigenetic and proteomic profiles, as well as performance in multiple functional bioassays.^16,18,33–36^ All cell-lines presented in this work possessed viability >95% (based on Trypan Blue exclusion assay) prior to plating for morphological profiling, immunosuppression assay, and secretomic profiling.

### 2.2 High Content Imaging, Morphological Profiling, and Morphological Landscapes

Morphological profiling was performed as described in ^19^ except modified to be performed in a high-throughput 96 well plate format. MSCs from each cell-line/passage experimental group were seeded at a density of 525 cells/cm^2^ in 96-well plates (Corning) and cultured for 24 hours in GM. GM was replaced with GM containing 64 different IFN-γ and TNF-α (Life Technologies) priming conditions consisting of a full factorial design of 0, 0.5, 1, 2, 5, 10, 20, 50 ng/mL of each cytokine (n=4 replicate wells for each cell-line/passage/priming condition) and cultured for an additional 24 hours. Following priming, MSCs were fixed with 4% paraformaldehyde for 15 minutes. Cell and nuclear morphology were assessed using 20 µM fluorescein-5-maleimide (Life Technologies) and 10 µg/mL Hoechst (Sigma-Aldrich), respectively. Samples for morphological analysis were imaged at 10X with a 6-by-6 stitched image captured for each well using an inverted Nikon Ti-S microscope with automated stage (Prior) and filters (Chroma Technology). Automated quantification of cellular and nuclear shape features was performed using CellProfiler v2.2.0^37^ (pipeline available in **File S1**) to obtain high dimensional single cell morphological data. An example of a segmented image output from the CellProfiler pipeline can be seen in **Figure S1**.

Morphological landscapes (i.e. 3D surface plots) were created for both single morphological features and a composite overall morphological score, which was created using principal component analysis (PCA). For each landscape plot (for a given cell-line/passage), the median of each single cell morphological feature for each well was averaged for quadruplicate wells and plotted for all 64 IFN-γ/TNF-α priming combinations to create a 3D surface plot using JMP Pro v14. An overall morphological landscape was plotted by first performing PCA on the high dimensional morphological data on a per well basis with each well consisting of 21-dimensional morphological data (definitions of each feature available in **Table 2**) selected based on features from our previous work.^19^ Principal component 1 (PC1) was taken to be a composite morphological score for each well and averaged across wells for all cell-lines/passages/priming conditions to create a 3D surface plot displaying morphology (PC1) versus [IFN-γ] versus [TNF-α].

### 2.3 Assessment of MSC T cell Suppression

We quantitatively assessed MSC suppression of activated T cells as described in our previous work using a MSC/PBMC (peripheral blood mononuclear cell) co-culture assay.^18,19^ For each experiment, MSCs from a given cell-line/passage were first seeded at 10,000 cells/well in 96-well plates (n=5 wells per cell-line/passage/priming condition) and cultured for 24 hours in GM. Then, GM was removed and the appropriate priming conditions were added to replicate wells. Following 24 hours of culture (unprimed and primed), 10^5^ PBMCs derived from a healthy human donor were stimulated with T cell activating beads (anti-CD3/CD28 Dynabeads, ThermoFisher) at a 1:1 PBMC:bead ratio in each well containing MSCs. Following 3 days of co-culture, PBMCs were collected and their activation assessed using flow cytometry (MACS-Quant, Miltenyi Biotec). Specifically, CD4^+^ and CD8^+^ T cells were individually assessed by proliferation (CFSE dilution), CD25 expression, and production of IFN-γ and TNF-α using FlowJo and compensation matrices generated using single-color control samples. All antibodies were purchased from BioLegend (San Diego, CA) and their information listed in **Table S3**.

### 2.4 MSC Secretomic Profiling

Quantitative analysis of MSC secreted proteins was performed using antibody arrays and ELISAs. For all secretion studies, MSCs from each cell-line at low passage were seeded in 12 well plates at a density of 10^4^ cells/cm^2^. Following 24 hours of culture in GM, the medium was removed and replaced with different priming conditions. After 24 hours of priming, the conditioned medium was collected, aliquoted into 1.5 mL microcentrifuge tubes, and stored at -80 C. Initially, we comprehensively profiled the secretome of one MSC line (PCBM1662 P3) using a 440-plex antibody array (Quantibody, Raybiotech, Norcross, GA). Frozen aliquots of conditioned medium and control GM (triplicate samples for each group) were shipped frozen to Raybiotech and analyzed using their array testing service, a Q440 Multiplex ELISA platform, which quantifies levels of 440 different human chemokines and cytokines. Secreted proteins found to significantly correlate with MSC T cell suppression for PCBM1662 P3 were then assessed for all cell-lines using ELISAs (Raybiotech) for the following target proteins: CXCL16, CXCL9, CXCL10, CXCL11, ICAM-1, CCL7, CCL8, CCL13, Legumain, Angiogenin (ANG), PLGF, DKK1. Secretion of Prostaglandin E2 (PGE2), a well-established MSC immunomodulatory factor^38^, was also quantified for each cell-line/priming condition using ELISA (Cayman Chemical, Ann Arbor, MI).

### 2.5 Statistical Analysis

All statistical tests were performed in GraphPad Prism v8 with the specific tests utilized for each experiment described in the figure legends.

## 3. RESULTS

### 3.1 MSCs exhibit greater morphological response to IFN-γ priming vs TNF-α priming

First, we assessed the effect of priming with either IFN-γ or TNF-α alone on MSC morphology. CellProfiler was used to measure 93 cell and nuclear morphological features of the MSCs. Selected morphological features for low passage MSCs are displayed in **Figure 1**. As little as 10 ng/mL of IFN-γ priming resulted in significant increases in cell major axis length, perimeter and cell aspect ratio after 24 hours compared to unprimed controls for nearly all MSC lines at low passage (8F3560 the exception for cell perimeter), although increasing this as high as 50 ng/mL did not have significant additional effects. On the other hand, cell solidity decreased with IFN-γ priming (p<0.05) for three of the MSC lines (8F3560 again the exception). TNF-α had a much less pronounced effect on morphological features with some distinct MSC line/passage responses e.g. decrease in cell aspect ratio and increase in cell solidity for RB9 with 50 ng/mL TNF-α priming. For high passage MSCs, IFN-γ again had a more pronounced effect than TNF-α with increased cell major axis length and aspect ratio for all cell-lines (**Figure S2**). Cell-line differences were observed in IFN-γ response for cell perimeter and solidity, and some TNF-α dependent responses were observed e.g. decrease in cell perimeter for 110877 and increase in cell solidity for RB9. Compared to day 0 (dotted lines, **Figure 1**), MSCs generally became larger (increased major axis length, perimeter), more elongated (increased aspect ratio), and more complex (decreased solidity) upon IFN-γ priming. For unstimulated and TNF-α only primed MSCs there was a notable decrease in major axis length and perimeter with some cell-line dependent differences observed in terms of aspect ratio or solidity when compared to day 0.

**Figure 1:**
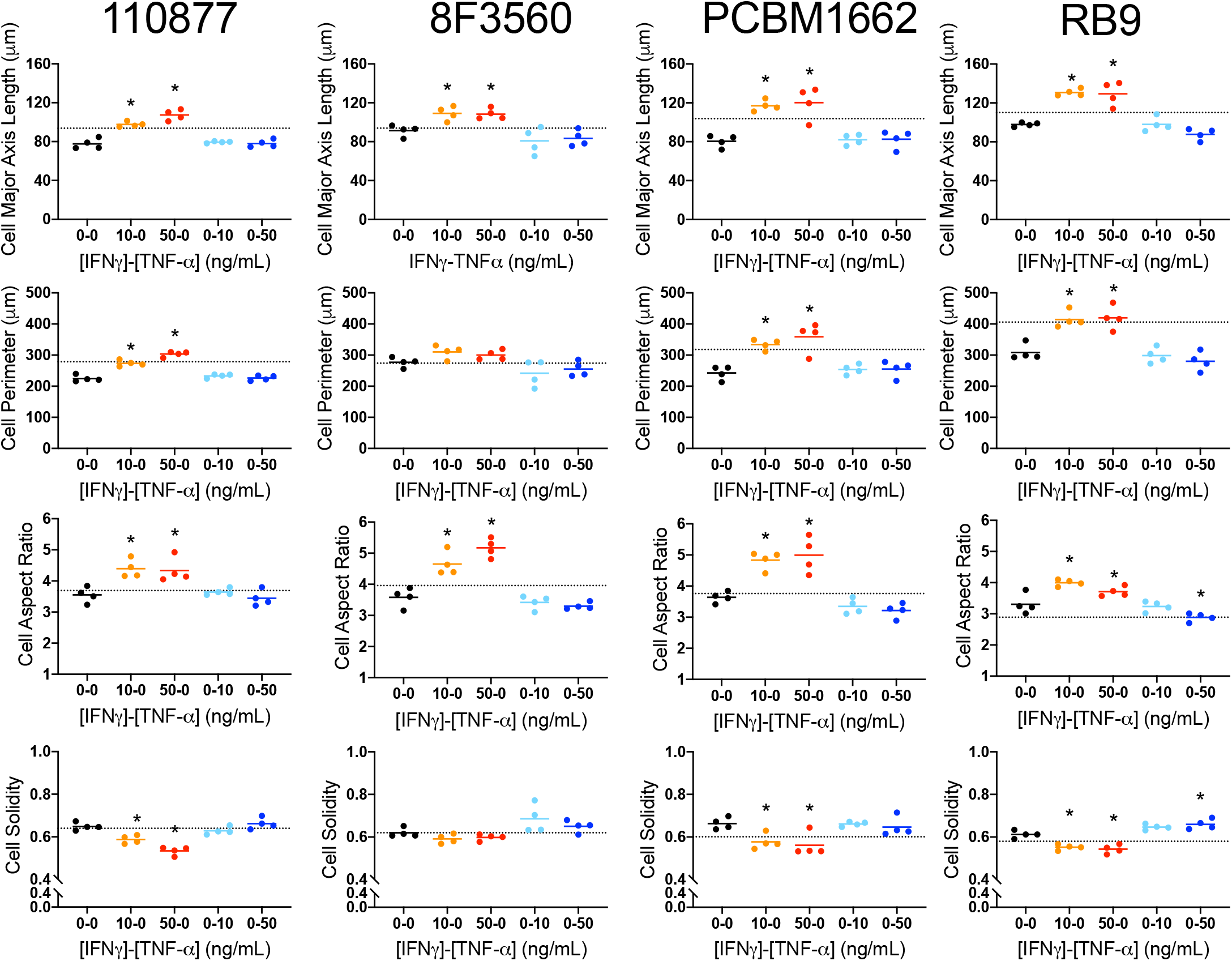
MSCs possess greater morphological response to IFN-γ only vs TNF-α only. Cell major axis length, cell perimeter, cell aspect ratio, and cell solidity of 4 MSC lines at low passage with different levels of IFN-γ and TNF-α priming. Reference (dotted) lines show day 0 values for each cell-line. One-way ANOVA with Dunnett’s multiple comparisons test vs unprimed control within the same cell-line. N = 4 wells per condition, *p < 0.05.

### 3.2 Synergistic effects of IFN-γ/TNF-α on MSC morphology

Next, we examined the combined effect of IFN-γ and TNF-α on MSC morphology. Early and late passage bone marrow-derived MSCs from four donors were primed with 64 different combinations of IFN-γ (0-50 ng/mL) and TNF-α (0-50 ng/mL) for 24 hours. Selected morphological features for low passage MSCs are displayed in **Figure 2**. We observed the same single factor response as in **Figure 1**, but with noticeable morphological responses occurring when MSCs were primed with both IFN-γ and TNF-α. For example, the major axis length and perimeter of the MSCs tended to increase with increasing IFN-γ concentration, with the addition of TNF-α contributing to a further increase. The inverse was true for cell form factor and solidity with both features decreasing significantly (with a threshold response observed at 5 ng/mL IFN-γ). While trends were consistent across MSC lines, the magnitude of the response varied considerably. For example, the perimeter of RB9 increased from ∼300 µm to ∼500 µm, compared to 110877, which increased from ∼250 µm to ∼350 µm. When comparing low versus high passage MSCs within a line, the morphological response (in terms of perimeter, for example) followed the same overall trend; however, cell-line dependent differences were observed due to different baseline/unstimulated morphologies for each cell-line (**Figure S3A)**. In all cases, the morphological response to priming was apparent at a threshold IFN-γ concentration of approximately 5 ng/mL and TNF-α concentration of 2 ng/mL. Increased priming past the threshold with higher concentrations of IFN-γ or TNF-α had a diminishing effect on morphological features.

**Figure 2:**
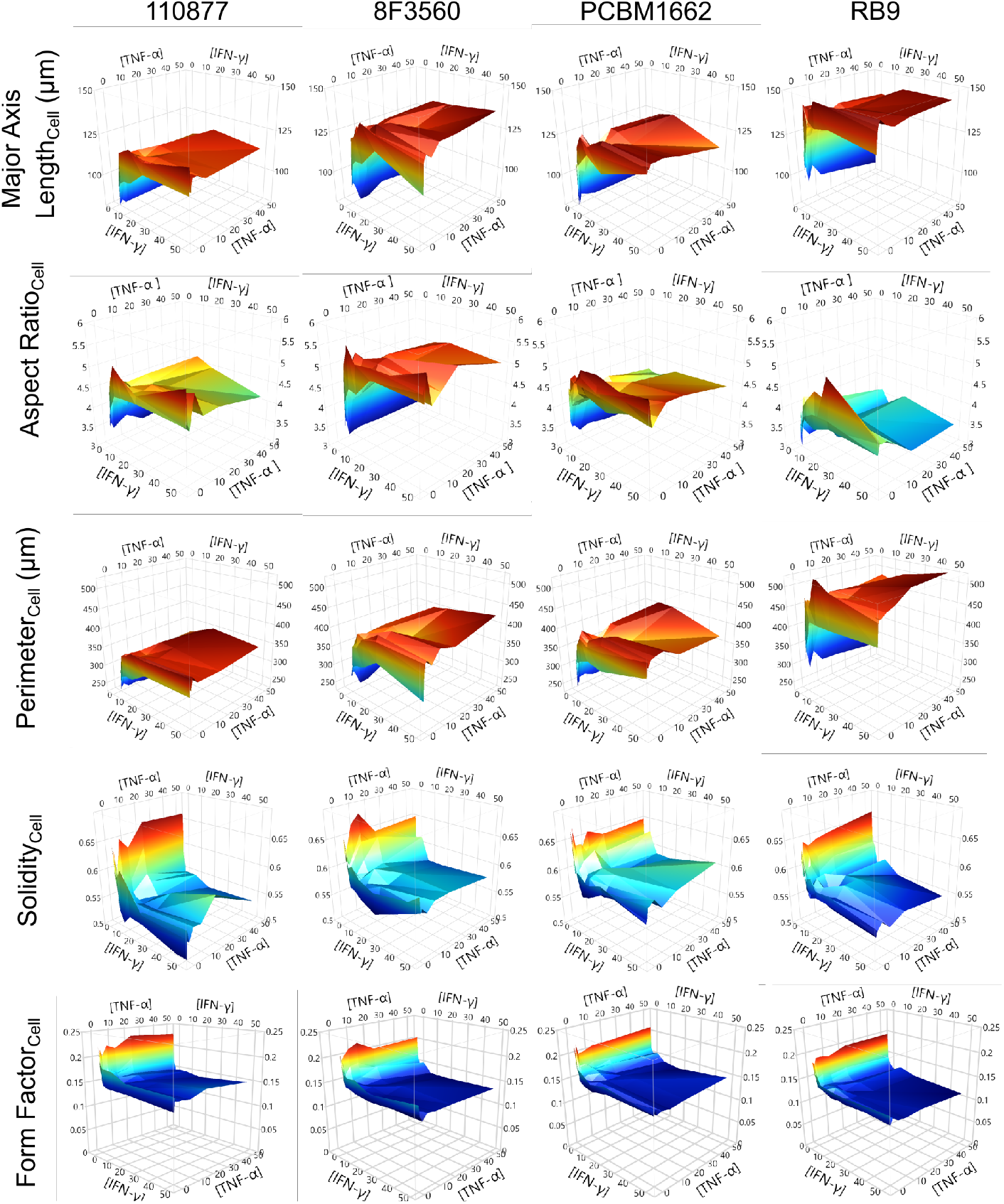
Synergistic effects of combined IFN-γ and TNF-α priming on MSC morphology. Average values of cell major axis length, aspect ratio, perimeter, solidity, and form factor of 4 MSC lines at low passage with different levels of IFN-γ and TNF-α priming. N = 4 wells for each IFN-γ/TNF-α priming condition with at least 200 cells analyzed per well.

Overall, IFN-γ priming resulted in the most significant change in morphology, with TNF-α contributing in a synergistic manner. Interestingly, in the case of cell solidity TNF-α appeared to ‘rescue’ MSCs from their IFN-γ mediated decrease (**Figure 2**). We also assessed MSC proliferation during the 24 hours of priming relative to their number prior to priming (**Figure S3B**,**C**). MSC proliferation followed a similar pattern to some of the morphological features in which it decreased with priming, the effect plateauing at higher concentrations. Like cell solidity, a similar ‘rescue effect’ was observed for proliferation as IFN-γ priming alone decreased cell number, but the addition of TNF-α mitigated this loss in cell number (compared to unprimed controls). Most MSC lines proliferated, although some high passage cell-lines had fewer cells after 24 hours.

### 3.3 Generation of a morphological landscape to visualize the overall morphological response

Given the high dimensionality of the morphological data and the fact that a single feature may not fully capture the effect of priming, we performed principal component analysis (PCA) on the morphological data to help visualize the overall MSC morphological response to priming (individual feature contributions to PCA shown in **Figure S4**). PCA was performed using the median value of 21 morphological features (**Table 2**) for each MSC line/passage/priming combination (512 data points). As the first principal component (PC1) generated from that analysis accounted for 62% of the variance in the data set, we used it as a composite measure of the overall MSC morphological response to priming (**Figure 3A, B**). Similar to the single morphological features, PC1 increased sharply with IFN-γ and TNF-α concentration before plateauing at higher concentrations. These changes were primarily in response to IFN-γ treatment, which were further augmented when combined with TNF-α concentrations at or above 2 ng/mL. Averaging PC1 across all MSC lines and passages revealed a distinct morphological landscape that effectively summarizes 21-dimensional morphological data from 4 cell-lines, two passages, and 64 different IFN-γ/TNF-α priming conditions (**Figure 3C**). In order to investigate the effect of priming on MSC function, we then chose 10 priming conditions based on key points within the morphological landscape to follow up on in future experiments (**Figure 3D**).

**Figure 3:**
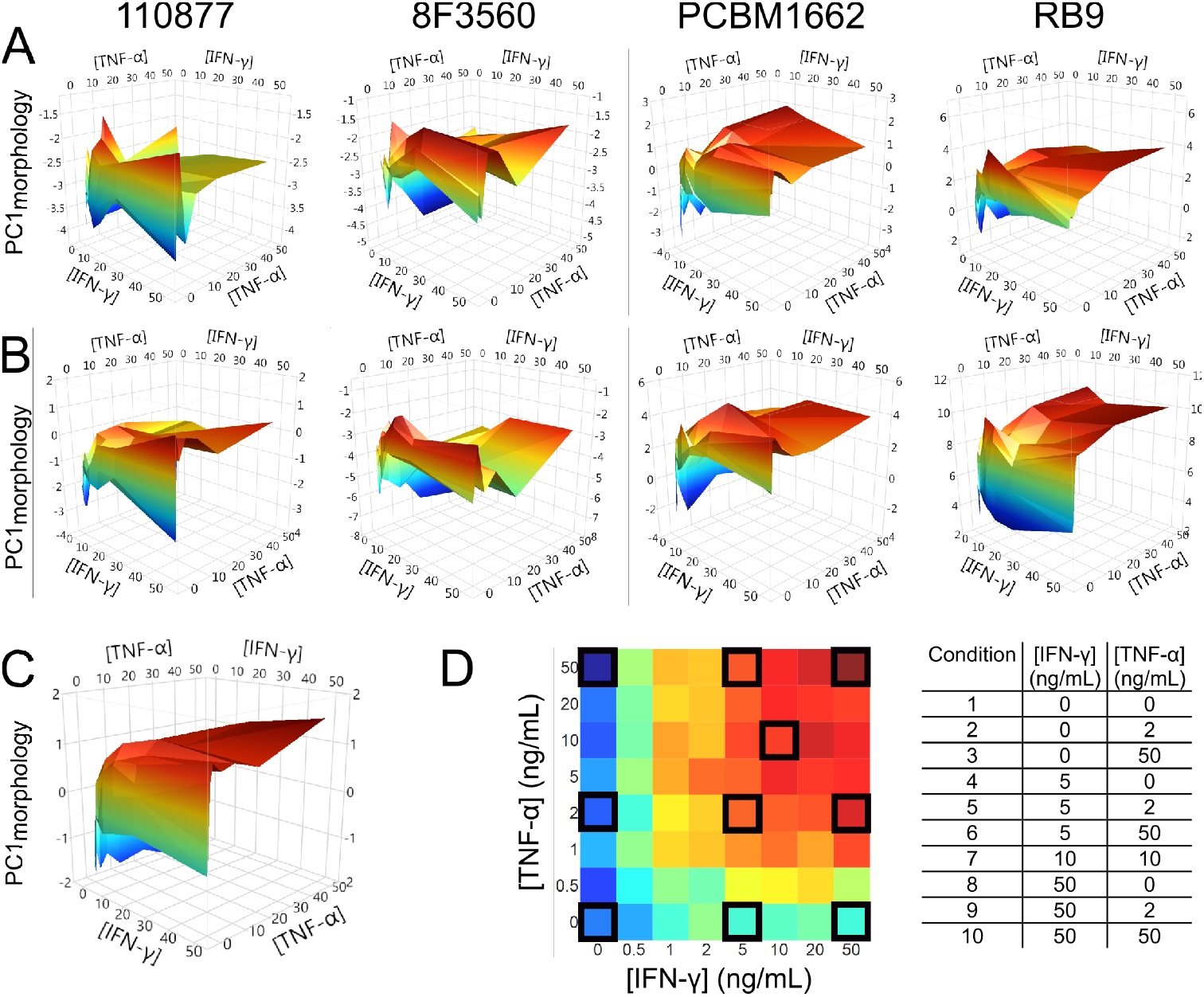
Morphological landscapes enable visualization of overall morphological response of MSCs to IFN-γ/TNF-α priming. (A) PC1_morphology_ vs [TNF-α] and [IFN-γ] at low passage. N = 4 wells per priming condition. (B) PC1_morphology_ vs [TNF-α] and [IFN-γ] at high passage. N = 4 wells per IFN-γ/TNF-α priming condition. (C) Average PC1_morphology_ vs [TNF-α] and [IFN-γ]. (D) Selected priming conditions (black boxes) for follow-up experiments summarized in tabular form.

### 3.4 Effects of combinatorial IFN-γ/TNF-α priming on T cell suppression

We then examined whether the different priming conditions identified from our screen had different/variable effects on the ability of MSCs to suppress activated T cells. T cell activation was measured via proliferation (%CFSE dilution), CD25 expression, TNF-α expression, and IFN-γ expression of CD4^+^ and CD8^+^ T cells that had been stimulated with anti-CD3/CD28 Dynabeads. Generally, T cell suppression was lower for all MSC lines at high passage (versus low passage) when MSCs were unprimed most notably for CD8^+^ T cell suppression (**Figure 4**). Although all of these activation parameters were affected by priming, the effect on proliferation was by far the most significant. While all MSCs suppressed CD4^+^ and CD8^+^ T cell proliferation, priming increased this suppression compared to that of unprimed MSCs.

**Figure 4:**
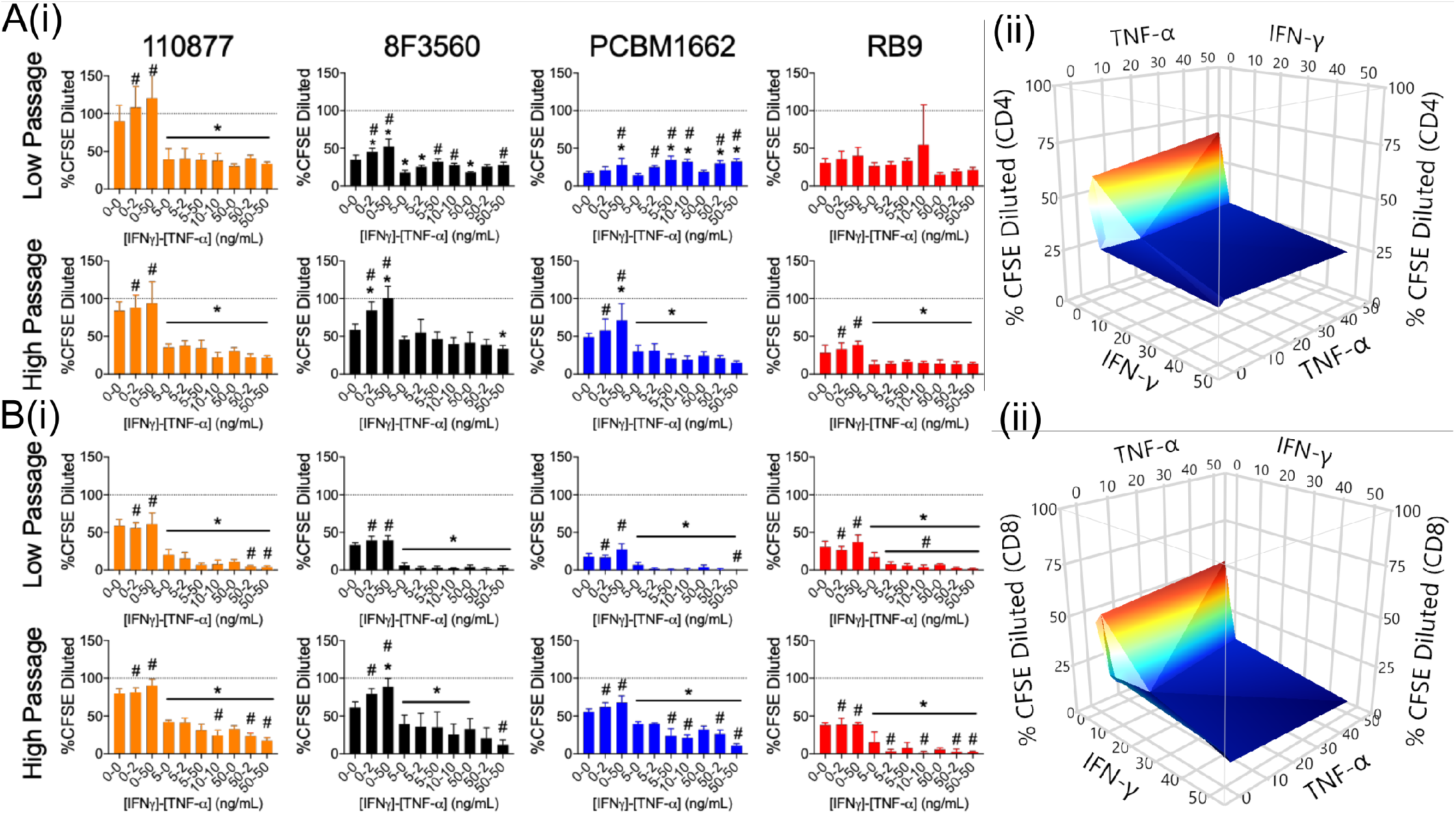
Effect of priming on MSC suppression of T cell proliferation. A(i) MSC suppression of CD4^+^ T cell proliferation as measured by CFSE dilution with TNF-α and IFN-γ priming by cell line and passage. N = 5 wells per priming condition. A(ii) MSC suppression of CD4^+^ and CD8^+^ T cell proliferation as measured by CFSE dilution vs [TNF-α] and [IFN-γ] averaged across all cell lines and passages. B(i) MSC suppression of CD8^+^ T cell proliferation as measured by CFSE dilution with TNF-α and IFN-γ priming by cell line and passage. N = 5 wells per priming condition. B(ii) MSC suppression of CD4^+^ and CD8^+^ T cell proliferation as measured by CFSE dilution vs [TNF-α] and [IFN-γ] averaged across all cell lines and passages. Reference dotted line represents activated PBMC-only control. Mean +/- SD. One-way ANOVA with Sidak’s multiple comparisons test. * denotes p < 0.05 vs unprimed control. # denotes p < 0.05 vs 5-0.

Similar to the morphological effects, MSC T cell suppression was primarily affected by IFN-γ priming, with effects becoming apparent at 5 ng/mL of IFN-γ only (**Figure 4**, *p<0.05). Conversely, TNF-α priming alone did not significantly affect MSC suppression of T cell activation. However, with combined IFN-γ and TNF-α priming, MSCs more effectively suppressed T cell proliferation than when primed by either cytokine alone (#p<0.05). This effect was particularly pronounced in the case of CD8^+^ T cell proliferation, in which as little as 5 ng/mL IFN-γ and 2 ng/mL TNF-α priming was sufficient to decrease proliferation significantly more than unprimed across all MSC lines and passages (p<0.05, **Figure 4B(i)**). The relationships between IFN-γ and TNF-α priming and MSC suppression of CD4^+^ and CD8^+^ T cell proliferation across all cell-lines/passages is summarized by **Figure 4A(ii)** and **Figure 4B(ii)**, respectively. Priming had less pronounced effects on CD25, TNF-α, and IFN-γ expression, although MSCs did generally suppress each of these measures (**Figures S5-S7**). Increased priming of MSCs also decreased the standard deviation of their suppression of both CD4^+^ and CD8^+^ T cell proliferation (i.e. decreasing functional heterogeneity) across MSC lines and passages (**Figure 5**) within a priming condition. The homogeneity in MSC function that resulted from maximal 50 ng/mL TNF-α + 50 ng/mL IFN-γ priming was remarkable considering the MSCs were from different donors/passages.

**Figure 5:**
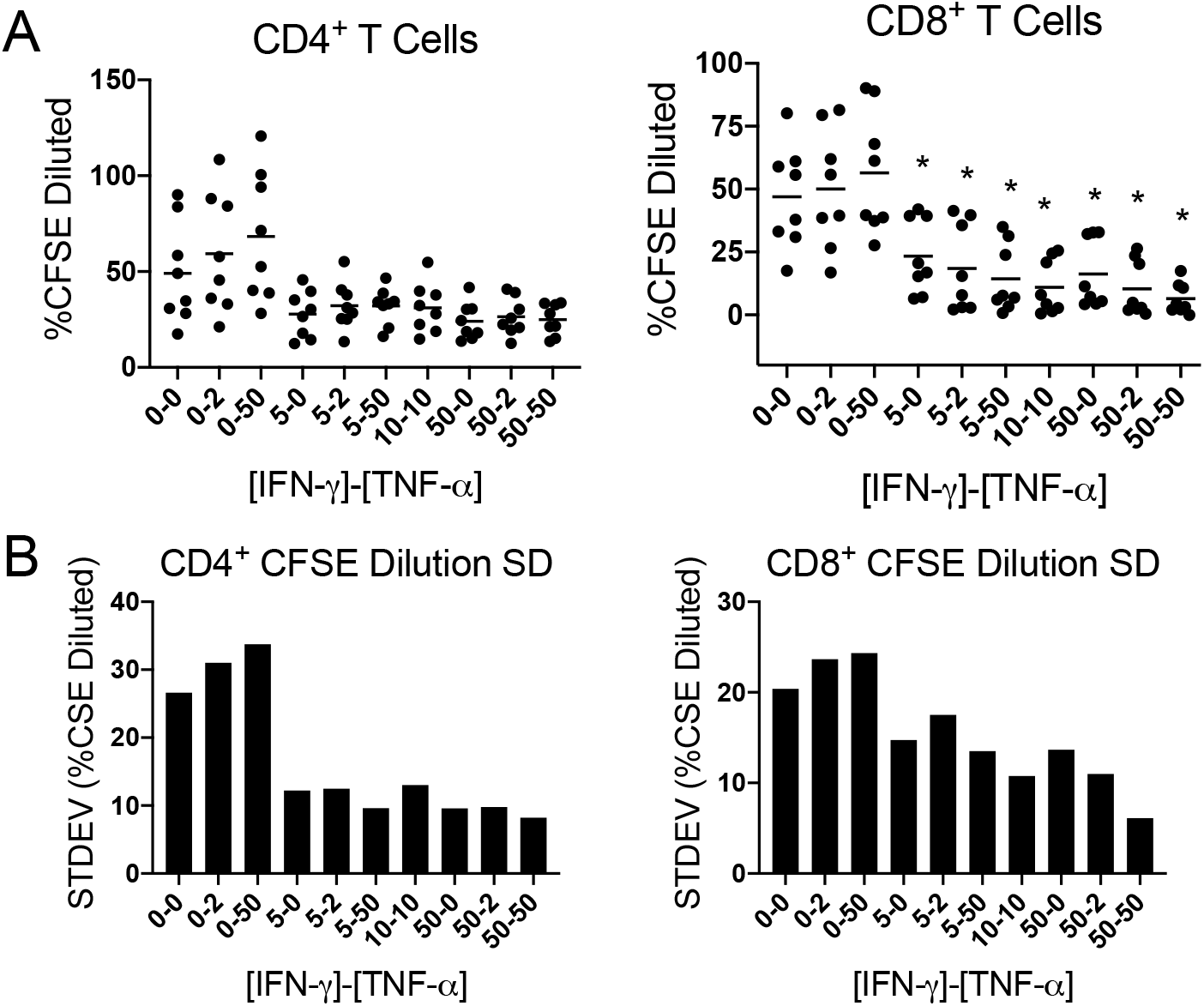
Decrease in MSC functional heterogeneity with priming. (A) Mean MSC suppression of CD4^+^ and CD8^+^ T cell proliferation as measured by CFSE dilution with TNF-α and IFN-γ priming across all cell lines and passages (N = 8). One-way ANOVA with Dunnett’s multiple comparisons post-hoc test * p < 0.05 different from unprimed control. (B) Standard deviations of mean MSC suppression of CD4^+^ and CD8^+^ T cell proliferation as measured by CFSE dilution with TNF-α and IFN-γ priming across all cell lines and passages.

### 3.5 MSC morphological response to priming is correlated with T cell suppression

As CD4^+^ and CD8^+^ T cell proliferation (as measured by CFSE dilution) had the most variance across priming conditions (**Table S4**), we used these functional metrics of T cell suppression to correlate with MSC morphological response to priming. Ten MSC morphological features were significantly correlated with function (Bonferonni-adjusted p-val cutoff < 3.44×10^−6^, **Table S5**). The two morphological features (cell form factor and nuclear major axis length) that exhibited the strongest correlations with T cell suppression are shown in **Figure 6**. As cell form factor decreased (increased priming), CD4^+^ and CD8^+^ T cell proliferation decreased (enhanced suppression). On the other hand, as the nuclear major axis length of MSCs increased (increased priming), proliferation of CD4^+^ and CD8^+^ T cells decreased (enhanced suppression). These correlations effectively encompass our above reported data for the effects of IFN-γ and TNF-α priming on MSC morphology and immunosuppressive function. The effects of varied priming conditions on MSC immunosuppression are mirrored by changes in many of their morphological features. However, there were some inconsistencies, notably PCBM1662 and RB9 at low passages but this could partially be explained by the fact that these cell-lines had high functional capacity when unstimulated and priming did not further enhance this function (**Figure 4A**, CD4^+^ T cells).

**Figure 6:**
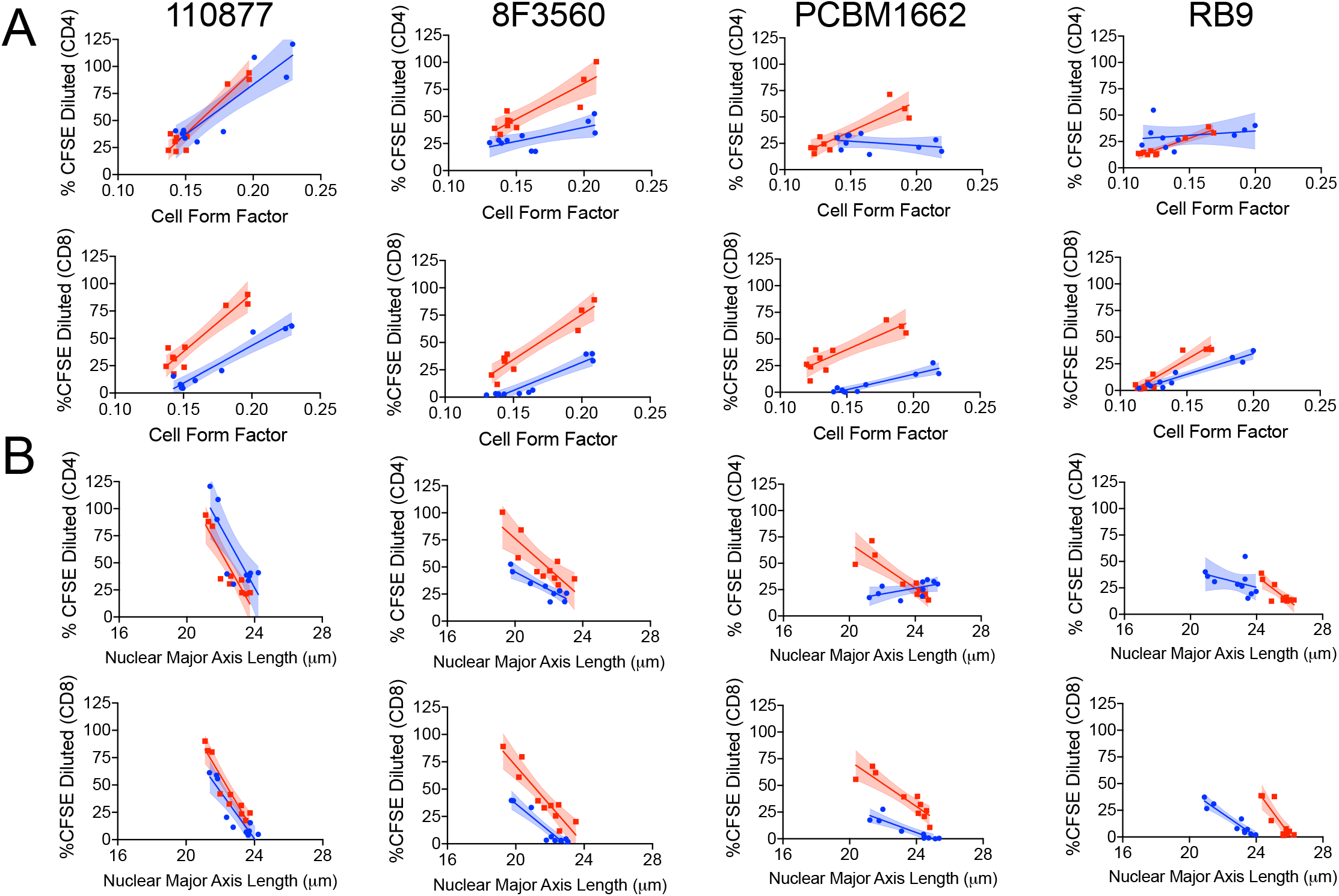
Morphological features correlate with MSC immunosuppression of activated T cells. (A) MSC suppression of CD4^+^ and CD8^+^ T cell proliferation vs MSC cell form factor by MSC line for low (blue) and high (red) passages. Simple linear regression, N = 10 priming conditions. (B) MSC suppression of CD4^+^ and CD8^+^ T cell proliferation vs MSC nuclear major axis length by MSC line for low and high passages. Simple linear regression, N = 10 priming conditions. Correlation coefficients for each graph are available in Table S5.

Overall, MSCs responded similarly between cell-lines and passages. In terms of morphology, MSCs became more spread and complex with priming, as shown by their increase in major axis length for both cell and nucleus, increased perimeter, and decreased solidity/form factor. This is in line with our previously reported results^18^ and has now been further demonstrated to be predictive of the effects of priming on MSC immunomodulation.

### 3.6 Response of MSC secretome to priming is correlated with T cell suppression

To better understand possible mechanisms of action for MSC-mediated T cell suppression, we profiled the secretome of MSCs primed with different combinations of IFN-γ/TNF-α. To identify target proteins, the conditioned media from one MSC line at low passage (PCBM1662 P3) was screened for 440 proteins after being primed using the 10 priming conditions identified from the morphological screen (**Figure 3D**). Proteins detected at concentrations both above the limit of detection and the media only control are listed in **Table S6**. Unsupervised hierarchical clustering performed on all priming conditions (**Figure 7A**) using a subset of proteins secreted at levels significantly higher than control medium (132 total) resulted in unprimed MSCs clustered at the top and maximally primed (50 ng/mL of both IFN-γ and TNF-α) clustered at the bottom of the heatmap. Following this, we correlated each secreted factor with T cell suppression for a given priming condition to determine whether any secreted factors could predict MSC function. 40 factors were significantly correlated with suppression in terms of CD8^+^ T cell proliferation (**Table S7**), which was again selected due to the highest observed CV (**Table S4**). From these, 12 of the factors with the highest correlation coefficients were selected as targets for performing follow-up ELISAs on all 4 MSC lines at low passage.

**Figure 7:**
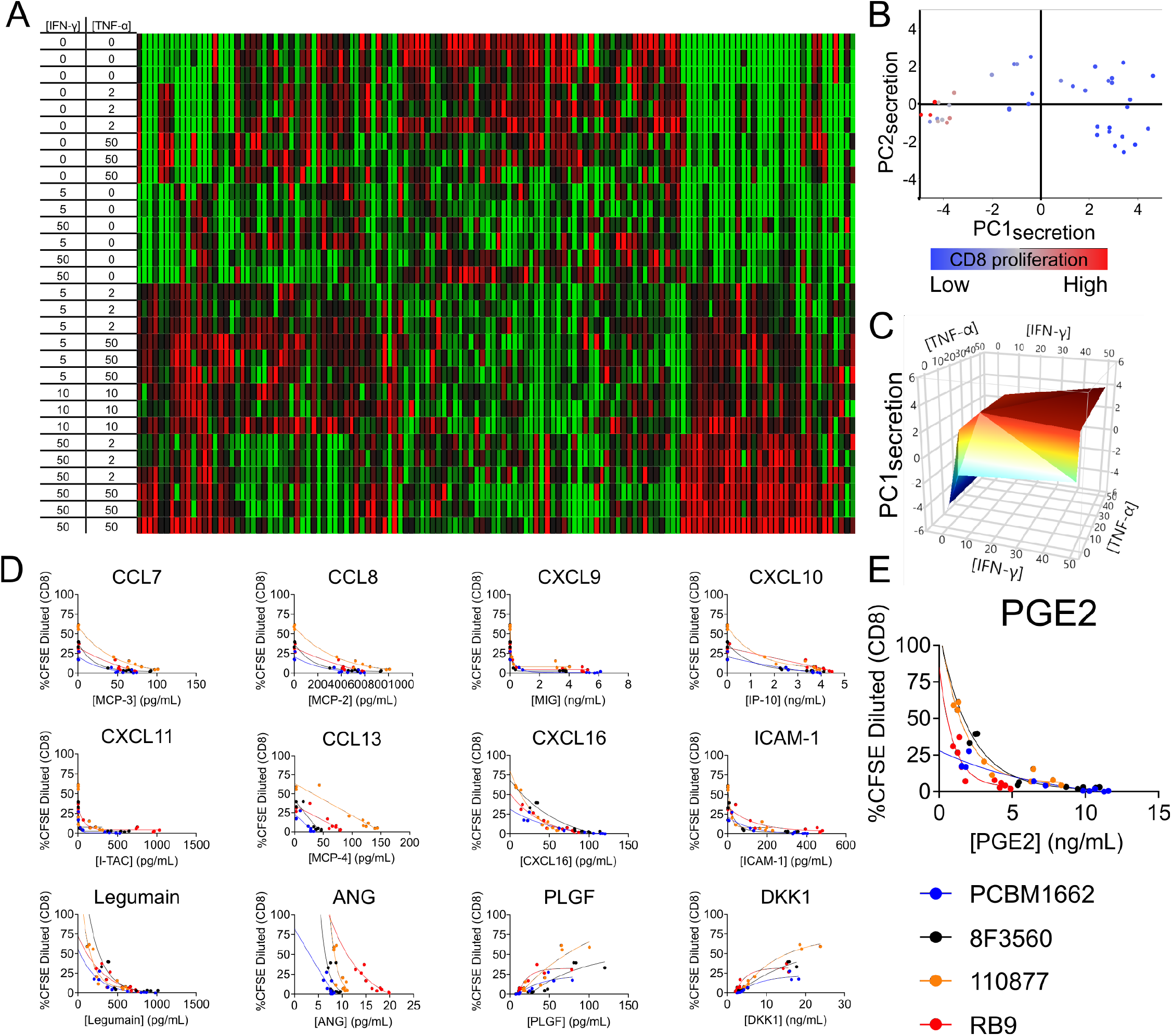
MSC secretome screening and correlation with T cell suppression. A) heat map of protein screen of PCBM1662 P3 conditioned media from 10 different priming conditions selected from morphological screen. N = 3 samples per priming condition. B) Principal Component Analysis of protein levels in MSC conditioned media from ELISA follow-up (12 proteins total) colored by CD8^+^ T cell proliferation. N = 40 (4 MSC lines x 10 priming conditions). C) PC1_secretion_ vs [TNF-α] and [IFN-γ]. N = 40. D) One phase decay, nonlinear fit of MSC suppression of CD8^+^ T cell proliferation vs levels of secreted proteins. N = 10 priming conditions per MSC line. E) One phase decay, nonlinear fit of MSC suppression of CD8^+^ T cell proliferation vs secreted PGE2. N = 10 priming conditions per MSC line.

Principal component analysis was performed on the secretory profiles generated from the ELISA follow-up (**Figure 7B**). Viewing the first and second principal components shows that PC1secretion tracks similarly with MSC function (colored by magnitude of CD8^+^ T cell proliferation). Plotting PC1secretion vs TNF-α and IFN-γ priming conditions reveals a similar landscape pattern (**Figure 7C**) to the morphological and functional landscapes shown in **Figure 3C** and **Figure 4**, respectively.

Furthermore, we correlated the secretion of these proteins with MSC suppression of CD8^+^ T cell proliferation and found strong relationships within MSC lines (**Figure 7D**). Most secreted proteins (CXCL9, CXCL10, CXCL11, CCL7, CCL8, CCL13, CXCL16, ICAM-1, Legumain, and ANG) increased with priming and were positively correlated with T cell suppression (lower %CFSE dilution). Conversely, PLGF and DKK1 were secreted at lower levels following priming and were thus negatively correlated with T cell suppression. The ELISA follow-up served to validate the initial secretome screen with additional cell-lines, as well as revealing some notable MSC line dependent correlations in the cases of CCL7, CCL8, CCL13, ANG, PLGF and DKK1. Secretion of other proteins, such as CXCL16, ICAM-1, CXCL9, CXCL10, CXCL11, and Legumain were shown to be MSC line independent in terms of their correlation with T cell suppression. Additionally, PGE2 (while not included in the initial screening array) was included in the ELISA follow-up due to its previously described involvement with MSC T cell suppression.^39^ It was found to correlate well with function independent of MSC line and increased with priming (**Figure 7E**).

## 4. DISCUSSION

This study explores in detail the response of MSCs to combinatorial TNF-α and IFN-γ priming and exemplifies the use of MSC morphology to screen for manufacturing conditions that enhance function. While inflammatory cytokine priming with TNF-α, IFN-γ (as well as other cytokines such as IL-1β) is a well-known method to improve MSC immunomodulatory capabilities, many of these studies have investigated only binary priming conditions (i.e. -/+ priming signals).^18,28,31,40,41^ Our work here encompasses four MSC lines (from three different commercial sources) at multiple passages and 10 priming conditions, with their functional effects (T cell suppression) being evaluated by eight different flow cytometry outcomes.

This comprehensive approach allowed us to identify a reduction of MSC heterogeneity across cell-lines and passages with priming. MSC functional heterogeneity can be attributed to a number of factors: different donor/tissue sources, extended culture, and manufacturing methods^10^. For example, a comparison of umbilical-, bone marrow-, and adipose tissue-derived MSCs found that adipose MSCs were better able to suppress the activation of PHA stimulated CD4^+^ and CD8^+^ T cells^42^. MSCs derived from the same tissue source but different donors can exhibit markedly different responses to IFN-γ as determined by production of the immunomodulatory enzyme IDO.^43^ Additionally, bone marrow-derived MSCs have been reported to have decreased secretion of immunomodulatory cytokines IL-6, IL-8, and RANTES with increased passage^44^. Similarly, our group has shown a decrease in the ability of MSCs to suppress the activation of CD4^+^ and CD8^+^ T cells with passaging.^18,19^ Functional heterogeneity of clonal MSC cultures has also been reported; however, much higher concentrations of combined IFN-γ and TNF-α priming were used to enhance immunomodulatory function and mitigate the observed heterogeneity^31^. The ability to reduce MSC heterogeneity – in terms of not only morphology, but also immunomodulatory function and secreted factors - opens potential new manufacturing approaches in which poorly performing MSC lines can be improved and their function effectively ‘rescued.’

This work provides a foundation for future studies to assess the effects of different priming conditions and predict MSC immunomodulation based on single cell responses. Previous studies attempting to screen MSC immunosuppressive function have utilized several approaches. Chinnadurai et al examined MSC secretome and RNA content as indicators of MSC suppression of CD3^+^ T cell proliferation.^15^ They found strong correlations between a number of cytokines in the media of PBMC-MSC cocultures and the observed MSC-mediated T cell suppression. Specifically, CXCL9 and CXCL10 were upregulated (both in terms of secreted protein levels and mRNA expression in cocultured and IFN-γ primed MSCs) and correlated with MSC suppression of T cells, which also was the case in our study. In another study, small molecules were screened for their ability to prime MSCs towards an immunosuppressive phenotype ^45^ using secretion of PGE2 as their target. Identified hits were followed up by examining how primed MSCs attenuated TNF-α secretion, first by macrophages *in vitro*, and finally in a mouse ear skin inflammation model. These approaches provide valuable information; however, they assess MSCs on a population level, while morphological profiling allows for single-cell resolution and potential identification of MSC functional subpopulations^46^. Additionally, relying on one functional measure (such as CD3^+^ T cell proliferation) or one analyte (such as PGE2) does not fully capture MSC multipotency i.e. their ability to modulate different immune cells and exert functions through multiple mechanisms of action^11^.

Here we demonstrated that priming MSCs with IFN-γ alone versus TNF-α alone induced a more significant response in terms of MSC morphology, T cell suppression, and secretion. Additionally, while TNF-α alone did not have a significant effect on MSC behavior, it did act synergistically with IFN-γ. Most studies involving MSC priming have examined the effects of either one cytokine or a combination of two cytokines, rather than a full factorial study examining different doses.^28^ The effects of binary priming MSCs with TNF-α and IFN-γ on immunomodulation was also investigated using MSCs from both bone marrow and Wharton’s jelly tissue sources.^47^ They found considerable heterogeneity between MSC donors and tissue sources in terms of the effect of TNF-α and IFN-γ priming on MSC suppression of PBMC proliferation, but did note that MSCs tended to become qualitatively larger and flatter with IFN-γ priming and more spindle-shaped upon exposure to TNF-α. This is consistent with our finding that MSC cell perimeter and major axis length increased with exposure to IFN-γ, but does not agree with our observations of TNF-α alone priming (**Figure 1**). In another study, Li et al reported that TNF-α priming had much greater effects than IFN-γ priming on MSC ability to suppress T cell proliferation, but also noted synergy when the two were used in combination ^48^. It is important to note that the priming in this referenced study was done concurrently with T cell coculture while priming in our study was done prior to coculture (i.e. pretreatment or preconditioning). Synergistic effects of 8 hour TNF-α/IFN-γ priming on MSC suppression of T cell proliferation has also been reported ^49^. Timing and duration of priming varies considerably between studies with demonstrated effects on MSC function, thus limiting the comparisons that can be made between studies and further emphasizing the critical need for standardization of MSC characterization assays.^29^

The MSC secretome has been implicated as the primary mechanism by which MSCs exert their immunomodulatory effects. Chinnadurai et al found significant correlations between MSC function as measured by cocultured T cell proliferation and their secretion of various cytokines and morphogens.^15^ Interestingly, they reported CXCL9 to have the same relationship with T cell proliferation i.e. reduced T cell proliferation/activation with increased secretion. CXCL9, CXCL10, CXCL11, CXCL16, and CCL8 are known chemokines for T cells^50–54^. Furthermore, CXCL9, CXCL10, and CXCL11 all bind CXCR3, which is primarily expressed on T cells and NK cells^50,52,55–58^. These chemokines may recruit activated immune cells to be locally modulated by other secreted or cell contact-mediated factors. CXCR3 is involved in regulatory T cell recruitment and migration as well, which could be another possible avenue for MSC immunomodulation.^59–61^

Secretion of DKK1, which inversely correlated with T cell suppression, is an inhibitor of canonical Wnt signalling^62^. PGE2, on the other hand, was positively correlated with MSC T cell suppression, and is a known activator of canonical Wnt signaling^63^. The Wnt pathway is a potent regulator of cell differentiation, growth, and migration, and its activation by primed MSCs may have far-ranging effects on the immune system. There is considerable evidence supporting PGE2 as an effector of MSC immunomodulation^64^ as it can suppress T cell activation and proliferation, and its secretion has been shown to be upregulated synergistically by IFN-γ/TNF-α primed MSCs, which is further supported by our results.^39,65^ Additionally, levels of PGE2 secretion have been shown to be predictive of MSC effects in a rat model of traumatic brain injury^38^. However, it is likely that MSCs exert their effects through multiple pathways and target cells, and the assessment of a single factor may not fully reflect MSC multipotency.^11^

The sensitivity, low cost and single-cell resolution of cell morphology could also be applied to many aspects of MSC manufacturing in order to detect and predict functional changes. For example, MSCs can respond to hypoxia or additional cytokines (e.g. IL-1β) through enhanced secretion of immunomodulatory factors and altered migratory capacity and therefore may exhibit distinct morphological responses to microenvironment signals besides IFN-γ and TNF-α.^66–68^ Additionally, our approach could be used to assess the impact of different manufacturing methods on MSC immunomodulation. Cell culture substrates and biomaterials can be tuned to direct MSC function and morphological profiling could be adapted to screen for biomaterial systems that further enhance MSC function.^69–72^ It is well established that MSCs lose function and become senescent over the course of *ex vivo* expansion and identification of soluble cues to include in defined growth medium could be another application of this morphology-based approach.^73,74^ Additionally, cryopreserved MSCs undergo a recovery period post-thaw, during which their function is impaired.^40^ Any observed differences in MSC immunosuppression caused by changes in these manufacturing methods could be assessed by the techniques described here.

Beyond screening for optimal MSC manufacturing conditions, morphological profiling could be used to assess manufacturing reagent batch-to-batch variability, which is an often overlooked challenge associated with cell manufacturing.^74–76^ Given that the MSC response to IFN-γ is consistent and predictable in this study (as well as in other studies), morphological profiling could be useful as a tool to assess the bioactivity of different batches of recombinant IFN-γ, TNF-α, or other priming factors. Culture medium used for MSC expansion represents an enormous source of variability when considering differences in supplement source (e.g. FBS vs. platelet lysate) and defined medium components (growth factors and small molecules).^74^ The effects of manufacturing changes on MSCs can be difficult to assess because there are no well-established CQAs associated with relevant MSC functions. This issue becomes costlier and more difficult to address as manufacturers advance in clinical development and often have to make significant changes (e.g. due to scaling or new vendor-sourced reagents) prior to performing Phase 3 studies and submitting a Biologics License Application. As we have demonstrated the ability of MSC morphology to predict reduction in functional heterogeneity with priming (**Figure 5**), we anticipate this approach could be similarly applied to reduce (and predict) functional heterogeneity derived from different media sources/compositions.

In summary, we have demonstrated that MSCs exhibit remarkably consistent morphological responses following IFN-γ priming that can be further enhanced, in a synergistic manner, with the addition of TNF-α. These morphological changes are strongly correlated with MSC immunosuppressive function, which in turn is reflected by secretion of chemokines and other immunomodulatory factors associated with T cell activation and migration. The morphological profiling approach presented in this work could be adapted and applied to improve manufacturing of other cell-types and cell-derived products (e.g. extracellular vesicles), and further explored as a means to better understand and control MSC functional heterogeneity.

## 5. Acknowledgments

The authors thank Drs. Zhaohui Ye, Saravanan Karumbayaram and Nirjal Bhattarai for their review of the manuscript, and Drs. Jessica Lo Surdo, Johnny Lam, and Eva Rudikoff for technical assistance in manufacturing the MSC lines used in this study. S.H.A. was supported by startup funds provided by the UGA College of Engineering to R.A.M. administered by the UGA Office of the Vice Provost of Research (OVPR). M.W.K. was supported in part by appointment to the Research Participation Program at the Center for Biologics Evaluation and Research (CBER) administered by the Oak Ridge Institute for Science and Education through the US Department of Energy and the US Food and Drug Administration (FDA). This work was also supported in part by the FDA Modernizing Science grant program, a Biomedical Advanced Research and Development Authority (BARDA) grant, and grant from the Medical Countermeasures Initiative and research funds from the Division of Cell and Gene Therapies.

## SUPPLEMENTAL FIGURES

**Figure S1.**
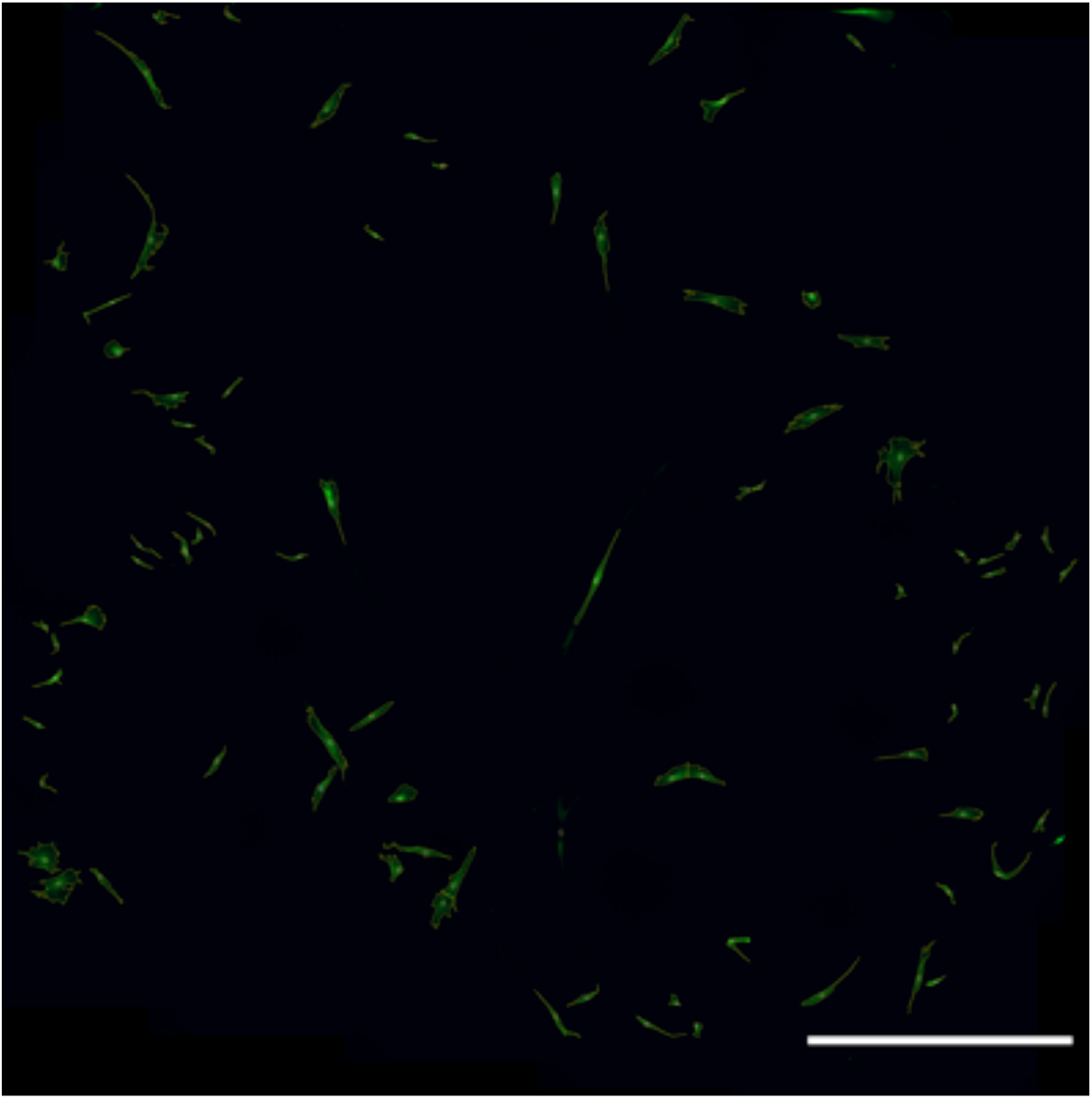
Representative large-field (6×6) output segmented image using custom CellProfiler pipeline (File S1). Scale bar = 1000 μm.

**Figure S2:**
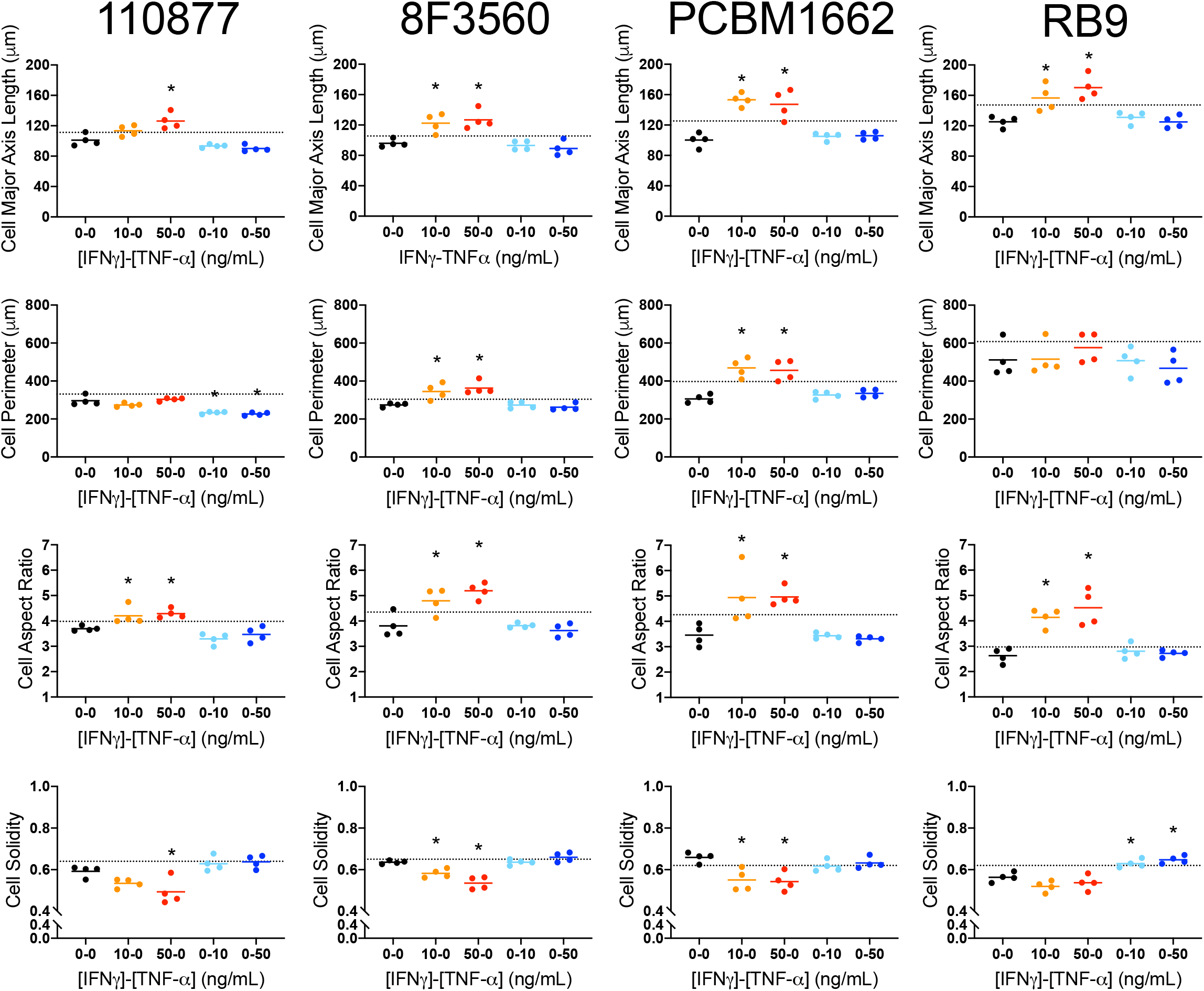
MSCs possess greater morphological response to IFN-γ only vs TNF-α only. Cell major axis length, cell perimeter, cell aspect ratio, and cell solidity of 4 MSC lines at low passage with different levels of IFN-γ and TNF-α priming. Reference (dotted) lines show day 0 values for each cell-line. One-way ANOVA with Dunnett’s multiple comparisons test vs unprimed control within the same cell-line. N = 4, *p < 0.05.

**Figure S3:**
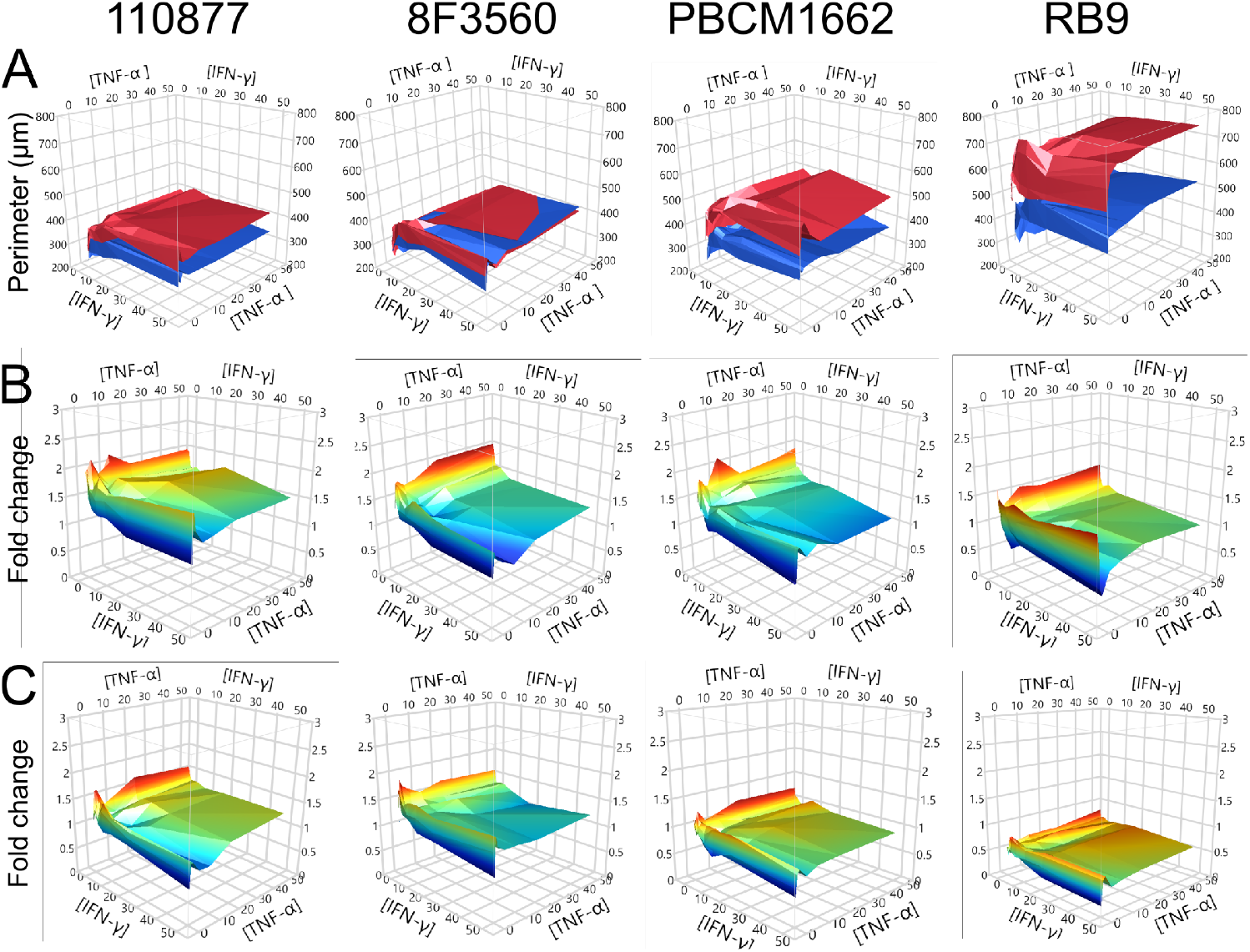
Effects of passage and interplay of IFN-γ and TNF-α on MSC growth and proliferation. (A) Average values of cell perimeter for low (blue) and high (red) passage MSCs across 4 MSC lines. N = 4 wells for all 64 priming conditions. (B) Average values of cell count fold change vs day 0 control for 4 MSC lines at low passage vs [TNF-α] and [IFN-γ]. N = 4 wells for all 64 priming conditions. (C) Average values of cell count fold change vs day 0 control for 4 MSC lines at high passage vs [TNF-α] and [IFN-γ]. N = 4 wells for all 64 priming conditions.

**Figure S4:**
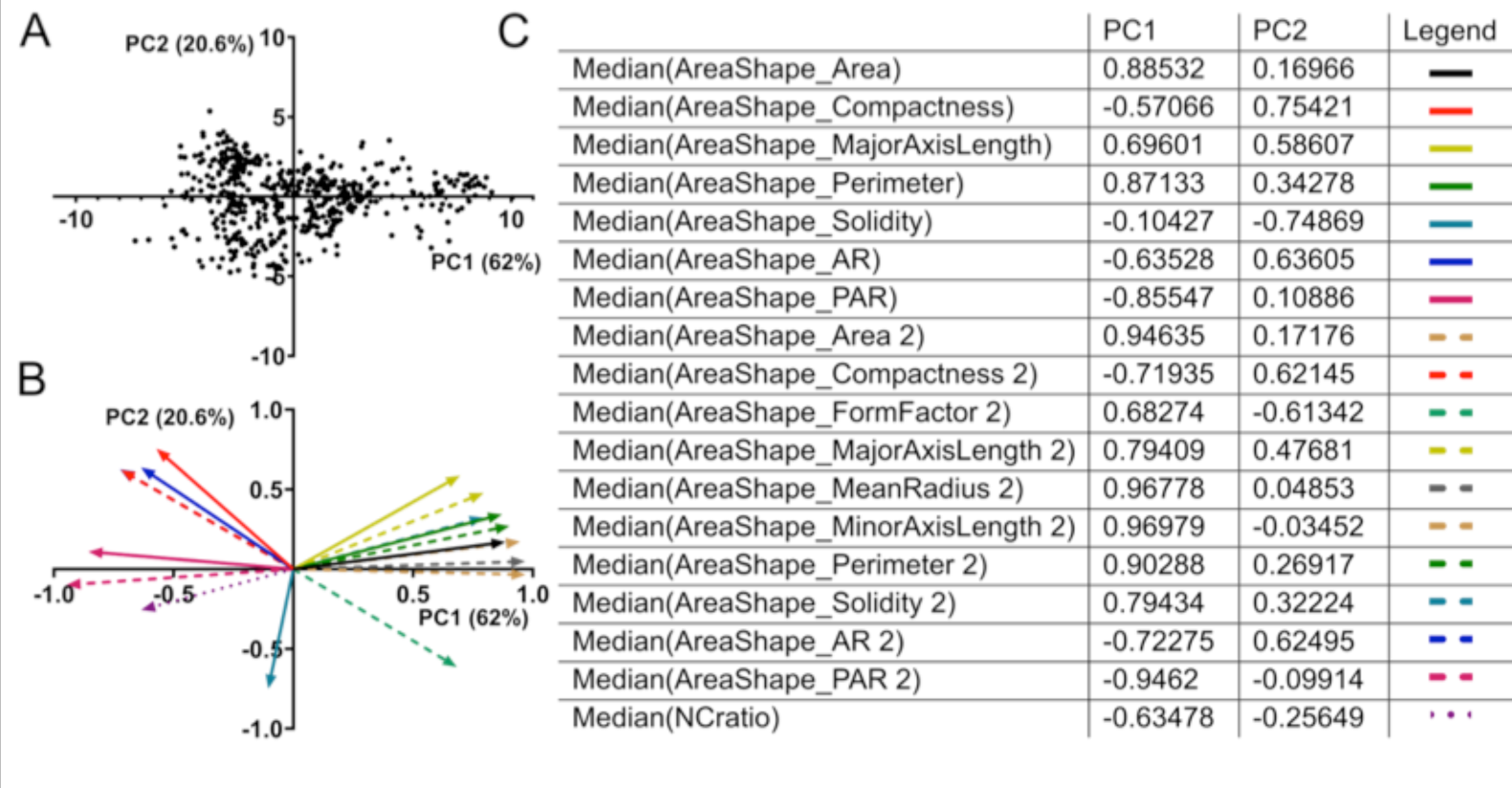
PCA of selected MSC morphological features. (A) Plot of PC1 vs PC2. 512 data points from 4 MSC lines, 2 passages, and 64 priming conditions. (B) Loading plot of morphological features with respect to PC1 and PC2. (C)

**Figure S5:**
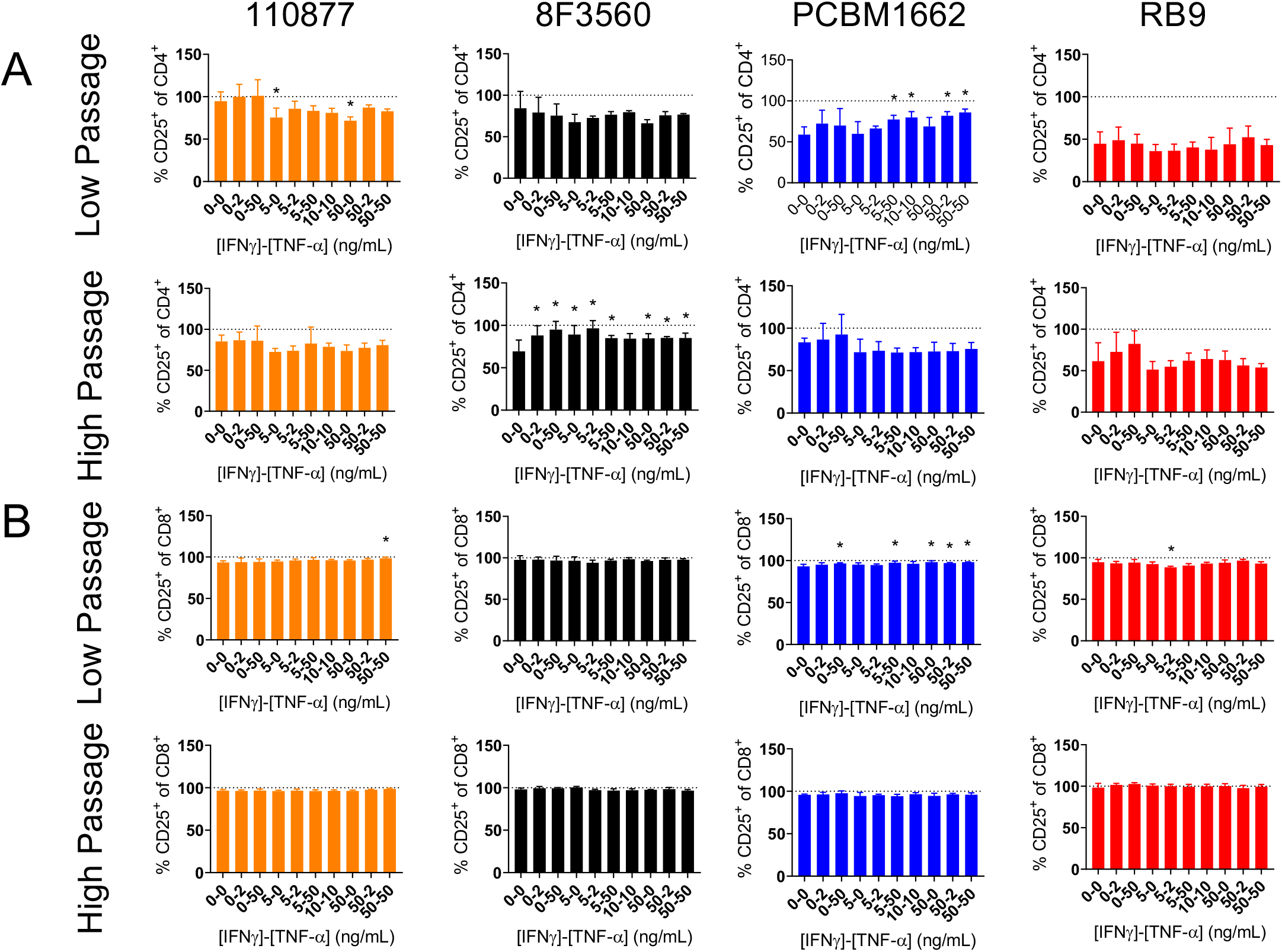
Effect of priming on MSC suppression of T cell CD25 expression. (A) MSC suppression of CD4^+^ T cell activation as measured by CD25 expression with TNF-α and IFN-γ priming by cell line and passage. (B) MSC suppression of CD8^+^ T cell activation as measured by CD25 expression with TNF-α and IFN-γ priming by cell line and passage. Reference line represents activated control PBMCs. Mean +/- SD. One-way ANOVA with Dunnett’s multiple comparisons test vs unprimed control. * denotes p < 0.05. N = 5 wells per priming condition.

**Figure S6:**
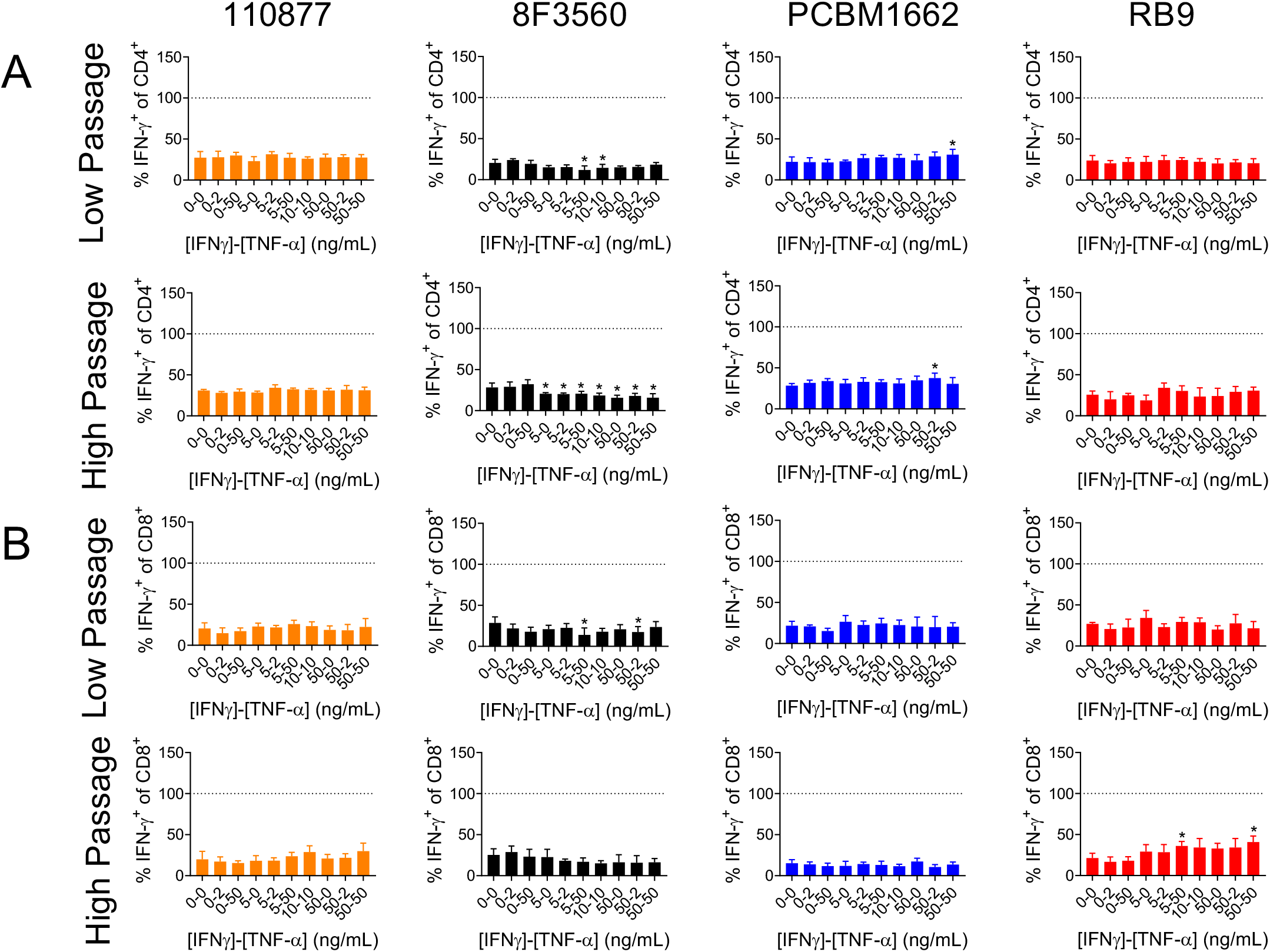
Effect of priming on MSC suppression of T cell IFN-γ expression. (A) MSC suppression of CD4^+^ T cell IFN-γ expression with TNF-α and IFN-γ priming by cell line and passage. (B) MSC suppression of CD8^+^ T cell IFN-γ expression with TNF-α and IFN-γ priming by cell line and passage. Reference line represents activated control PBMCs. Mean +/- SD. One-way ANOVA with Dunnett’s multiple comparisons test vs unprimed control. * denotes p < 0.05. N = 5 wells per priming condition.

**Figure S7:**
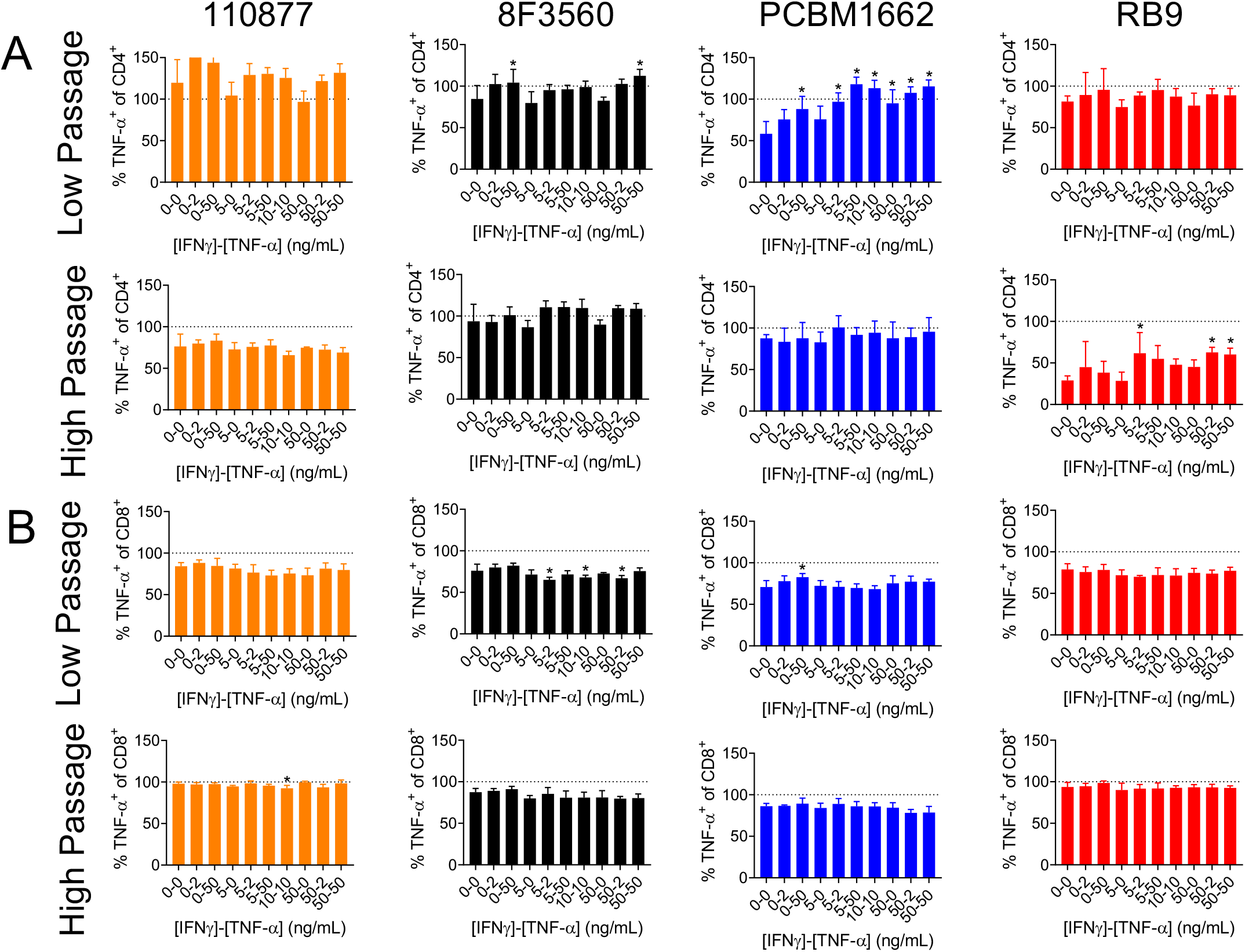
Effect of priming on MSC suppression of T cell TNF-α expression. (A) MSC suppression of CD4^+^ T cell TNF-α expression with TNF-α and IFN-γ priming by cell line and passage. (B) MSC suppression of CD8^+^ T cell TNF-α expression with TNF-α and IFN-γ priming by cell line and passage. Reference line represents activated control PBMCs. Mean +/- SD. One-way ANOVA with Dunnett’s multiple comparisons test vs unprimed control. * denotes p < 0.05. N = 5 wells per priming condition.

## SUPPLEMENTAL TABLES

**Table S1.**
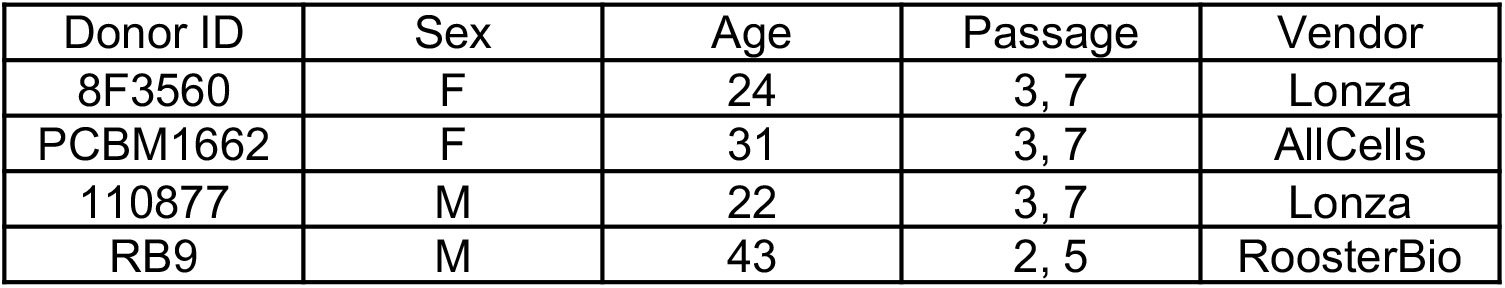
Donor information for all MSC lines used in this study.

| Donor ID | Sex | Age | Passage | Vendor |
| --- | --- | --- | --- | --- |
| 8F3560 | F | 24 | 3, 7 | Lonza |
| PCBM1662 | F | 31 | 3, 7 | AllCells |
| 110877 | M | 22 | 3, 7 | Lonza |
| RB9 | M | 43 | 2, 5 | RoosterBio |

**Table S2.** Definitions for morphological features used in this study. All morphological features are evaluated using CellProfiler’s MeasureObjectSizeShape module except for Aspect Ratio, Perimeter/Area, and Nuclear/Cytoplasm Ratio (NC Ratio).

|  | Morphological Feature | Units | Feature Definition |
| --- | --- | --- | --- |
| Cellular Features | Area | $\mu\text{m}^2$ | Total pixels within the object |
|  | Aspect Ratio | dimensionless | Major Axis Length divided by Minor Axis Length |
|  | Compactness | dimensionless | Variance of the radial distance of the object's pixels from the centroid divided by the area |
| | Form Factor | dimensionless | $4 \cdot \pi \cdot \text{Area} / \text{Perimeter}^2$ |
| | Major Axis Length | $\mu\text{m}$ | The length (in pixels) of the major axis on the ellipse that has the same normalized second central moments as the region |
| | Mean Radius | $\mu\text{m}$ | The mean distance of any pixel in the object to the closest pixel outside the object |
| | Minor Axis Length | $\mu\text{m}$ | The length (in pixels) of the minor axis on the ellipse that has the same normalized second central moments as the region |
| | Perimeter/Area | $\mu\text{m}^{-1}$ | Perimeter of the object divided by the area of the object |
| | Perimeter | $\mu\text{m}$ | The total number of pixels around the boundary of each the object |
|  | Solidity | dimensionless | The proportion of the pixels in the convex hull that are also in the object |
| Nuclear Features | Area | $\mu\text{m}^2$ | Total pixels within the object |
|  | Aspect Ratio | dimensionless | Major Axis Length divided by Minor Axis Length |
|  | Compactness | dimensionless | Variance of the radial distance of the object's pixels from the centroid divided by the area |
| | Form Factor | dimensionless | $4 \cdot \pi \cdot \text{Area} / \text{Perimeter}^2$ |
| | Major Axis Length | $\mu\text{m}$ | The length (in pixels) of the major axis on the ellipse that has the same normalized second central moments as the region |
| | Mean Radius | $\mu\text{m}$ | The mean distance of any pixel in the object to the closest pixel outside the object |
| | Minor Axis Length | $\mu\text{m}$ | The length (in pixels) of the minor axis on the ellipse that has the same normalized second central moments as the region |
| | Perimeter/Area | $\mu\text{m}^{-1}$ | Perimeter of the object divided by the area of the object |
| | Perimeter | $\mu\text{m}$ | The total number of pixels around the boundary of each the object |
|  | Solidity | dimensionless | The proportion of the pixels in the convex hull that are also in the object |
|  | NC Ratio | dimensionless | Area of the nucleus divided by the area of the cell |

**Table S3.** Antibodies used for flow cytometric assessment of T cell activation.

| Antigen | Clone | Conjugation | Manufacturer/catalog no. | Validation Profile |
| --- | --- | --- | --- | --- |
| hCD4 | RPA-T4 | PerCP | BioLegend 300528 | 1DegreeBio |
| hCD8a | RPA-T8 | APC | BioLegend 301049 | 1DegreeBio |
| hCD25 | M-A251 | Pacific Blue | BioLegend 356130 | Antibodypedia |
| hIFN-γ | B27 | PE/Cy7 | BioLegend 506518 | 1DegreeBio |
| hTNF-α | Mab11 | Pacific Blue | BioLegend 502920 | 1DegreeBio |

**Table S4:**
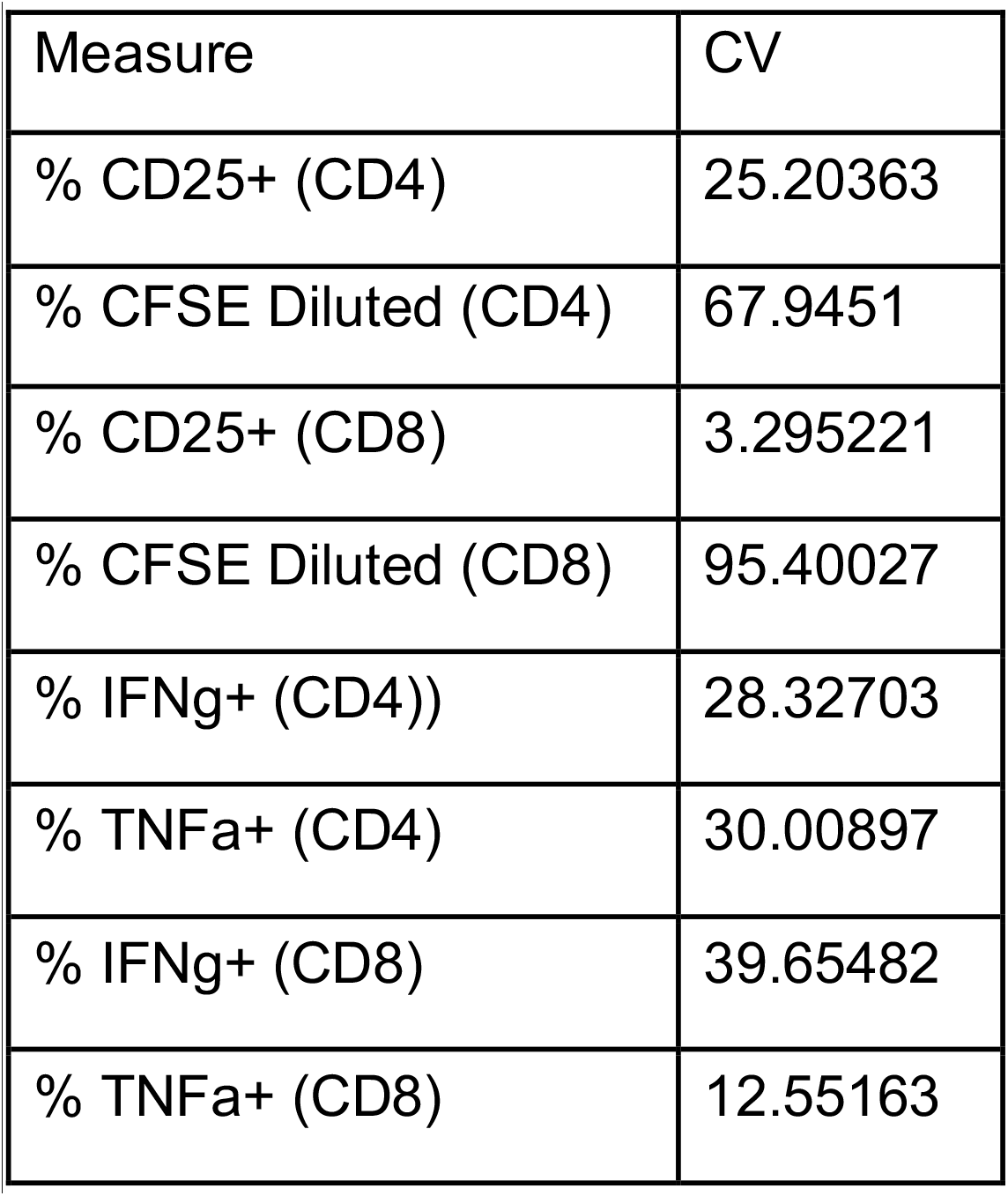
MSC suppression of T cell proliferation varies with MSC priming. Coefficient of variance for measures of T cell activation across all MSC lines/passages and priming conditions (80 total).

**Table S5:** MSC morphological features correlate with their suppression of T cell proliferation. Significantly correlated MSC morphological/functional pairs. N = 10. Bonferonni-adjusted p value cutoff calculated as p< 3.44×10^−6^ (0.05/14537 tests)

| Functional measure | Morphological Feature | Correlation | Signif Prob |
| --- | --- | --- | --- |
| % CFSE Diluted (CD4) | 24cell_FormFactor | 0.6489 | 7.53E-11 |
| % CFSE Diluted (CD8) | 24cell_FormFactor | 0.6419 | 1.40E-10 |
| % CFSE Diluted (CD4) | 24cell_MajorAxisLength | -0.6277 | 4.62E-10 |
| % CFSE Diluted (CD8) | 24cell_MajorAxisLength | -0.5513 | 1.16E-07 |
| % CFSE Diluted (CD4) | 24cell_Perimeter | -0.5666 | 4.29E-08 |
| % CFSE Diluted (CD4) | 24nuc_Area | -0.5863 | 1.10E-08 |
| % CFSE Diluted (CD8) | 24nuc_Area | -0.5368 | 2.85E-07 |
| % CFSE Diluted (CD4) | 24nuc_MajorAxisLength | -0.6096 | 1.96E-09 |
| % CFSE Diluted (CD8) | 24nuc_MajorAxisLength | -0.6494 | 7.22E-11 |
| % CFSE Diluted (CD4) | 24nuc_MeanRadius | -0.5667 | 4.25E-08 |
| % CFSE Diluted (CD4) | 24nuc_MinorAxisLength | -0.5344 | 3.29E-07 |
| % CFSE Diluted (CD4) | 24nuc_PerimAreaRatio | 0.6023 | 3.41E-09 |
| % CFSE Diluted (CD8) | 24nuc_PerimAreaRatio | 0.5361 | 2.98E-07 |
| % CFSE Diluted (CD4) | 24nuc_Perimeter | -0.6029 | 3.25E-09 |
| % CFSE Diluted (CD8) | 24nuc_Perimeter | -0.5893 | 8.91E-09 |
| % CFSE Diluted (CD4) | 24nuc_Solidity | -0.5047 | 1.81E-06 |
| % CFSE Diluted (CD8) | 24nuc_Solidity | -0.5625 | 5.61E-08 |

**Table S6:**
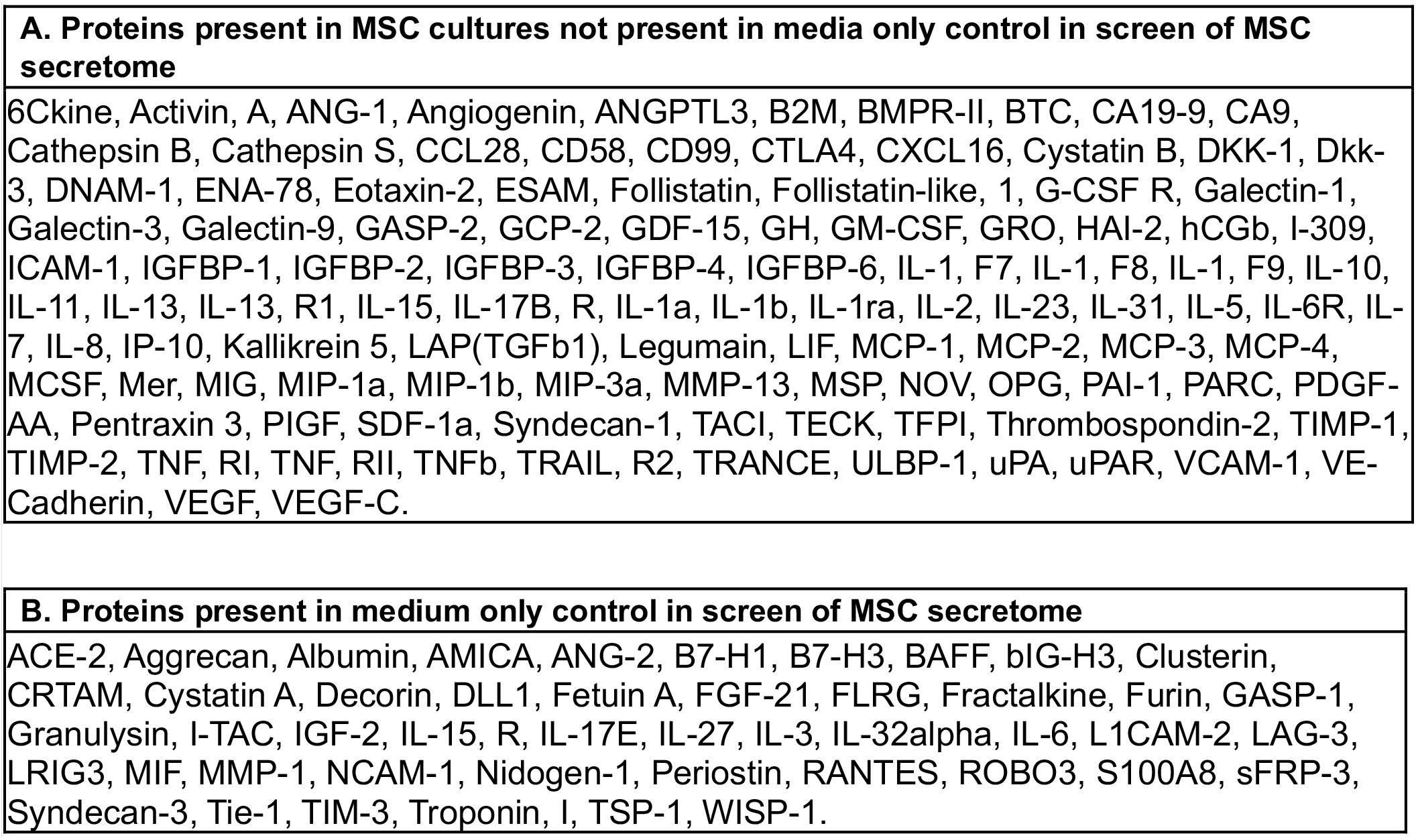
List of proteins present in MSC cultures and control MSC growth medium. (A) proteins detected in at least one conditioned medium sample from a single MSC line/priming condition that were also not detected in medium-only controls. (B) proteins present in MSC growth medium only controls

**Table S7:** Correlation of MSC secretome with immunosuppression. Correlations of T cell proliferation with specific secreted factors identified from the secretome profiling screen for low passage MSC line PCBM1662 ordered by p-value (low to high) determined through linear regression. Bonferonni-corrected cutoff p-value=0.05/132 = 3.8×10^−4^.

| Variable | by Variable | Correlation | Signif Prob |
| --- | --- | --- | --- |
| %CFSE Diluted (CD8) | MCP-2 | -0.9224 | 4.32E-13 |
| %CFSE Diluted (CD8) | IFNg | -0.9104 | 3.03E-12 |
| %CFSE Diluted (CD8) | IP-10 | -0.9029 | 8.87E-12 |
| %CFSE Diluted (CD8) | DKK-1 | 0.8965 | 2.08E-11 |
| %CFSE Diluted (CD8) | PIGF | 0.883 | 1.07E-10 |
| %CFSE Diluted (CD8) | CXCL16 | -0.8323 | 1.19E-08 |
| %CFSE Diluted (CD8) | MCP-3 | -0.819 | 3.16E-08 |
| %CFSE Diluted (CD8) | Legumain | -0.8087 | 6.42E-08 |
| %CFSE Diluted (CD8) | ANG-1 | 0.7922 | 1.83E-07 |
| %CFSE Diluted (CD8) | MCP-4 | -0.7898 | 2.11E-07 |
| %CFSE Diluted (CD8) | I-TAC | -0.7878 | 2.38E-07 |
| %CFSE Diluted (CD8) | ICAM-1 | -0.7874 | 2.44E-07 |
| %CFSE Diluted (CD8) | ANGPTL3 | 0.7711 | 6.13E-07 |
| %CFSE Diluted (CD8) | SDF-1a | 0.7671 | 7.63E-07 |
| %CFSE Diluted (CD8) | MIG | -0.7668 | 7.73E-07 |
| %CFSE Diluted (CD8) | B2M | -0.7577 | 1.25E-06 |
| %CFSE Diluted (CD8) | MCSF | -0.7509 | 1.75E-06 |
| %CFSE Diluted (CD8) | TIMP-2 | 0.7298 | 4.73E-06 |
| %CFSE Diluted (CD8) | CTLA4 | 0.724 | 6.12E-06 |
| %CFSE Diluted (CD8) | CA9 | 0.724 | 6.12E-06 |
| %CFSE Diluted (CD8) | IL-1ra | -0.7222 | 6.62E-06 |
| %CFSE Diluted (CD8) | IL-11 | -0.7218 | 6.74E-06 |
| %CFSE Diluted (CD8) | IL-2 | -0.7035 | 1.45E-05 |
| %CFSE Diluted (CD8) | Cathepsin S | -0.7027 | 1.50E-05 |
| %CFSE Diluted (CD8) | IL-1 F9 | 0.697 | 1.87E-05 |
| %CFSE Diluted (CD8) | TNFb | -0.6903 | 2.43E-05 |
| %CFSE Diluted (CD8) | CD99 | 0.6888 | 2.57E-05 |
| %CFSE Diluted (CD8) | IL-13 | -0.6847 | 3.00E-05 |
| %CFSE Diluted (CD8) | G-CSF R | 0.6791 | 3.69E-05 |
| %CFSE Diluted (CD8) | IL-5 | -0.6777 | 3.88E-05 |
| %CFSE Diluted (CD8) | VEGF | 0.6655 | 5.99E-05 |
| %CFSE Diluted (CD8) | CA19-9 | 0.6622 | 6.72E-05 |
| %CFSE Diluted (CD8) | IL-7 | -0.6615 | 6.89E-05 |
| %CFSE Diluted (CD8) | IL-6 | -0.6593 | 7.42E-05 |
| %CFSE Diluted (CD8) | CCL28 | -0.6543 | 8.78E-05 |
| %CFSE Diluted (CD8) | RANTES | -0.6533 | 9.07E-05 |
| %CFSE Diluted (CD8) | GCP-2 | 0.648 | 1.08E-04 |
| %CFSE Diluted (CD8) | IL-1a | -0.646 | 1.15E-04 |
| %CFSE Diluted (CD8) | ENA-78 | 0.6454 | 1.17E-04 |
| %CFSE Diluted (CD8) | TSP-1 | 0.6304 | 1.88E-04 |

